# *Cis*-aconitate therapy protects against influenza mortality by dual targeting of viral polymerase and ERK/AKT/NF-κB signaling

**DOI:** 10.1101/2025.06.18.660279

**Authors:** Adeline Cezard, Déborah Brea-Diakite, Virginie Vasseur, Alan Wacquiez, Loic Gonzalez, Ronan Le Goffic, Bruno Da Costa, Delphine Fouquenet, Severine Heumel, Arnaud Machelart, Eik Hoffmann, Priscille Brodin, François Trottein, Cyrille Mathieu, Lola Canus, Florentine Jacolin, Pierre-Olivier Vidalain, Laure Perrin-Cocon, Vincent Lotteau, Julien Burlaud-Gaillard, Dominique Tertigas, Michael G. Surette, Antoine Legras, Damien Sizaret, Thomas Baranek, Christophe Paget, Antoine Guillon, Mustapha Si-Tahar

## Abstract

Influenza virus poses a significant global health challenge, causing approximately 500,000 deaths annually. Its ability to evade antiviral treatments and vaccine-induced immunity underscores the need for novel therapeutic approaches. Our study identifies *cis*-aconitate (*cis*-aco), a mitochondria-derived metabolite, as a potent dual-action agent against influenza, independently of its metabolic derivative, itaconate. *Cis*-aco impairs viral polymerase activity, suppressing viral mRNA expression and protein synthesis to inhibit replication across a range of influenza subtypes. This antiviral efficacy is confirmed in *ex vivo* human airway and lung organotypic models. Beyond its antiviral properties, *cis-aco* exhibits potent anti-inflammatory effects, disrupting key inflammatory cascades and reducing the secretion of inflammatory mediators. In a mouse model of influenza pneumonia, *cis*-aco mitigates viral replication, inflammation, and immune cell activation, significantly improving survival. Notably, its efficacy persists even when administered at later stages of infection, when oseltamivir/Tamiflu® is no longer effective. These findings position *cis*-aco as a promising influenza treatment, combining antiviral and anti-inflammatory benefits within a clinically relevant timeframe.

## INTRODUCTION

Influenza viruses have long been major causes of morbidity and mortality, with heightened attention since the 1918 pandemic, driving extensive research into therapies [1]. Current approaches, including vaccination and antivirals, often demonstrate limited effectiveness. The short duration of vaccine-induced immunity, combined with the intrinsic antigenic drift of influenza viruses, undermines sustained protection [2, 3]. Skepticism also persists regarding the efficacy of approved anti-influenza drugs, especially when administered later in the course of infection [4–6]. Therefore, developing innovative strategies effective even after infection onset is crucial.

The pathophysiology of influenza-related pneumonia stems from the intrinsic viral pathogenicity and the immune response. While a robust immune response is essential for viral clearance, excessive cellular recruitment and the release of cytotoxic molecules can lead to lung hyper-inflammation, resulting in tissue damage, morbidity, and death [7, 8].

Interestingly, the recent discovery of metabolic reprogramming of immune cells has opened avenues for innovative therapeutic approaches [9–13]. We and others have demonstrated that hosts develop metabolic countermeasures in response to infection. [9, 14–19]. For instance, using an integrated approach combining metabolomics, *in vitro*, and *in vivo* infection assays, we recently discovered the inhibitory effect of tricarboxylic acid cycle (TCA)-derived succinate on influenza virus infection. This inhibition is primarily associated with succinylation and nuclear retention of the viral nucleoprotein, although additional mechanisms may also contribute [16].

Building on this finding, our investigation explored a broader range of host metabolites for their ability to regulate influenza virus infection in human lung epithelial cells. Among the TCA intermediates examined, *cis*-aconitate (*cis*-aco) stood out for its potent antiviral and anti-inflammatory properties. We further elucidated the molecular mechanisms underlying *cis*-aco’s protective effect, and our *in vivo* experiments confirmed its efficacy in mitigating severe influenza pneumonia. Remarkably, *cis*-aco demonstrated therapeutic benefits even when administered at advanced stages of infection, when conventional treatment with oseltamivir (Tamiflu®, [20]) proved ineffective.

## RESULTS

### *Cis-*aco inhibits both viral replication and production of inflammatory mediators in influenza virus-infected lung epithelial cells

Building on our previous work examining the anti-influenza effects of succinate [16], we selected eight metabolites derived from glycolysis or the TCA cycle for evaluation. Human bronchial epithelial BEAS-2B cells were infected with influenza A virus (IAV; A/Scotland/20/74, H3N2) and treated with 3.4 mM of each metabolite at 4 hours (h) post-infection (p.i.) (Fig. 1a). This concentration was chosen as the highest dose without cytotoxic effects across all tested metabolites (Fig. EV1a-b). Notably, *cis*-aco treatment at this concentration had no significant impact on cell proliferation, mitochondrial mass, or ROS production (Fig. EV1c-e).

**Figure 1.**
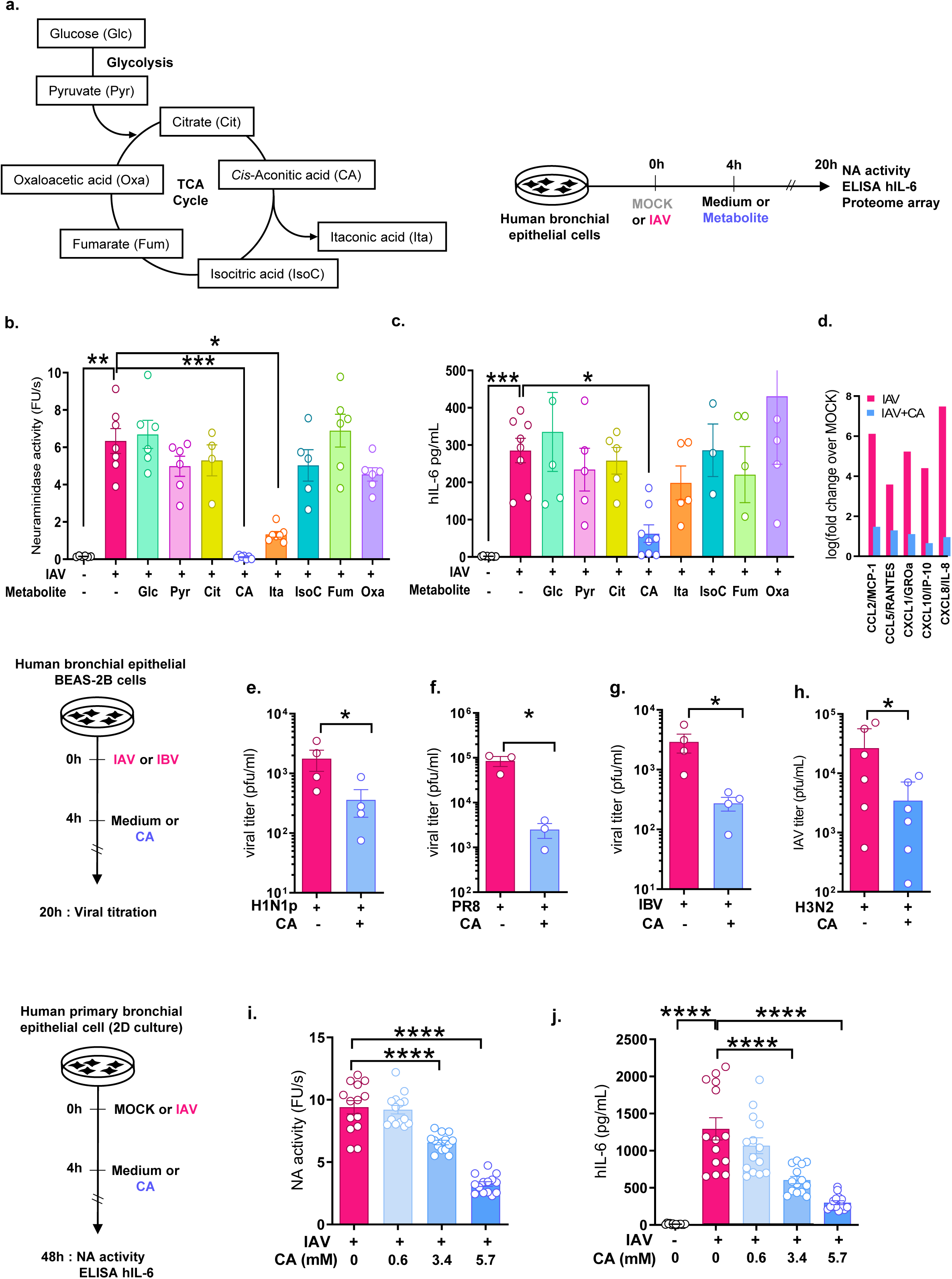
Anti-influenza properties of *cis*-aco among glycolysis and TCA cycle metabolites. **(a-d)** Human bronchial epithelial (BEAS-2B) cells were infected with influenza A/Scotland/20/74 (H3N2) virus at MOI=1 (IAV) or left uninfected (MOCK) for 4 h, then treated (Metabolite) or untreated (Medium) with 3.4 mM of TCA cycle or glycolysis metabolites for 16h. **(a)** Metabolites tested: (*cis*-aco (CA), itaconic acid (Ita), oxaloacetic acid (Oxa), isocitric acid (IsoC), citrate (Cit), fumarate (Fum), pyruvate (Pyr) or glucose (Glc)). **(b)** Viral particle production assessed by neuraminidase activity assay. (**c)** hIL-6 levels in cell supernatants measured by ELISA. **(d)** Immune mediator levels in cell supernatants determined using a protein-array. Pink bars represent fold changes in mediator secretion induced by IAV infection, and blue bars represent fold changes in IAV + CA conditions relative to MOCK. **(e-h)** BEAS-2B cells were infected with: **(e)** influenza A/pandemic/2009 H1N1 (H1N1p) strain, **(f)** A/Puerto Rico/8/1934 H1N1 (PR8) virus, **(g)**influenza B Yamagata (B/Paris/234/2013) virus (IBV) or **(h)** A/Scotland/20/74 (H3N2) strain. At 4 hours p.i., cells were washed and treated or not for 16 h with 3.4 mM of *cis*-aco (CA). Production of infectious viral particles in cell supernatants was quantified by plaque-forming unit assay. **(i-j)** PBEC in two-dimensional (2D) liquid culture were infected with influenza A/Scotland/20/74 (H3N2) virus at MOI=1 for 4 h, then treated or not with varying concentrations of *cis*-aco (CA) for 44 h. At 48 h p.i., neuraminidase activity **(i)** and hIL-6 levels **(j)** were measured in cell supernatants to assess viral particle production and pro-inflammatory cytokine release. Data are presented as the mean ± SEM. Results represent cumulative data from 1 **(d)**, 7 **(b)**, 3 **(g),** or 4 **(c, e, f, h, i, j)** independent experiments. PBEC data **(i-j)** are shown as the mean ± SEM from duplicate samples of PBEC from 4 independent individuals. Statistical analyses were performed using the Kruskal-Wallis test with Dunn’s multiple comparison test or one-way ANOVA (**i-j)**.

To assess the release of neo-virions, we measured neuraminidase (NA) activity in cell supernatants at 20 hours p.i.. Treatment with *cis*-aco resulted in a 60-fold reduction in NA activity (p=0.0003), while itaconate treatment led to a smaller, yet significant, 4.8-fold reduction (p=0.028) (Fig. 1b). No significant changes in NA activity were observed with other metabolites. Concomitantly, *cis*-aco treatment also reduced the levels of the inflammatory cytokine interleukin-6 (IL-6) in IAV-infected epithelial cells (p=0.03) (Fig. 1c). We next evaluated the expression of a broader panel of pro-inflammatory cytokines and chemokines, including CCL2/MCP-1, CCL5/RANTES, CXCL1/GROα, CXCL10/IP-10, IL-6, and CXCL8/IL-8. In IAV-infected cells, the levels of these mediators were increased by 4-to 7-fold compared to MOCK conditions (Fig. 1d, pink bars). *Cis*-aco treatment inhibited this inflammatory response by at least 3-fold (Fig. 1d, blue bars). Of note, *trans*-aconitate exhibited similar antiviral and anti-inflammatory activities (Fig. EV2), suggesting that both *cis*– and *trans*-isomers of aconitate possess anti-influenza properties.

Both influenza A and B viruses contribute to seasonal flu epidemics, with influenza A encompassing a diverse range of subtypes [21]. To assess the broad-spectrum antiviral potential of *cis*-aco, we evaluated its activity against several influenza strains: influenza A/pandemic/2009 H1N1 (Fig. 1e), influenza A/Puerto Rico/8/1934 H1N1 (Fig. 1f), influenza B Yamagata strain B/Paris/234/2013 (Fig. 1g), in comparison with our “reference” strain influenza A/Scotland/20/74 H3N2 (Fig. 1h). Plaque-forming unit assays revealed a significant decrease in viral particle production in cells treated with *cis*-aco, irrespective of influenza virus type or subtype (p<0.05; Fig. 3a-c).

The foregoing results were obtained using the immortalized bronchial epithelial BEAS-2B cell line, which can exhibit altered functional and metabolic profiles [22, 23]. To better reflect the lung’s physiological conditions, we next tested *cis*-aco in two dimensional (2D) cultures of primary bronchial epithelial cells. In these cells, *cis*-aco treatment significantly reduced IAV release in a dose-dependent manner (p<0.0001; Fig. 1i) and inhibited IAV-induced IL-6 production by up to 80% (Fig. 1j). *Cis*-aco also decreased IAV-induced cell death by approximately 70%, as shown by SYTOX™ labeling (Fig. EV3). While 2D primary bronchial epithelial cultures are closer to *in vivo* conditions than immortalized cell lines, they still lack the complexity of native tissues. To address this, we developed a 3D culture model using primary airway epithelial cells grown at an air-liquid interface (ALI), forming a differentiated, polarized epithelium with basal, goblet, and ciliated cells (Fig. 2a). This model exhibited a transepithelial electrical resistance of ∼400 ohms/cm² and showed beating cilia (not shown). In this more complex system, *cis*-aco again demonstrated anti-viral and anti-inflammatory effects, confirming its efficacy in a model that more closely mimics the *in vivo* environment (Fig. 2b-d).

**Figure 2.**
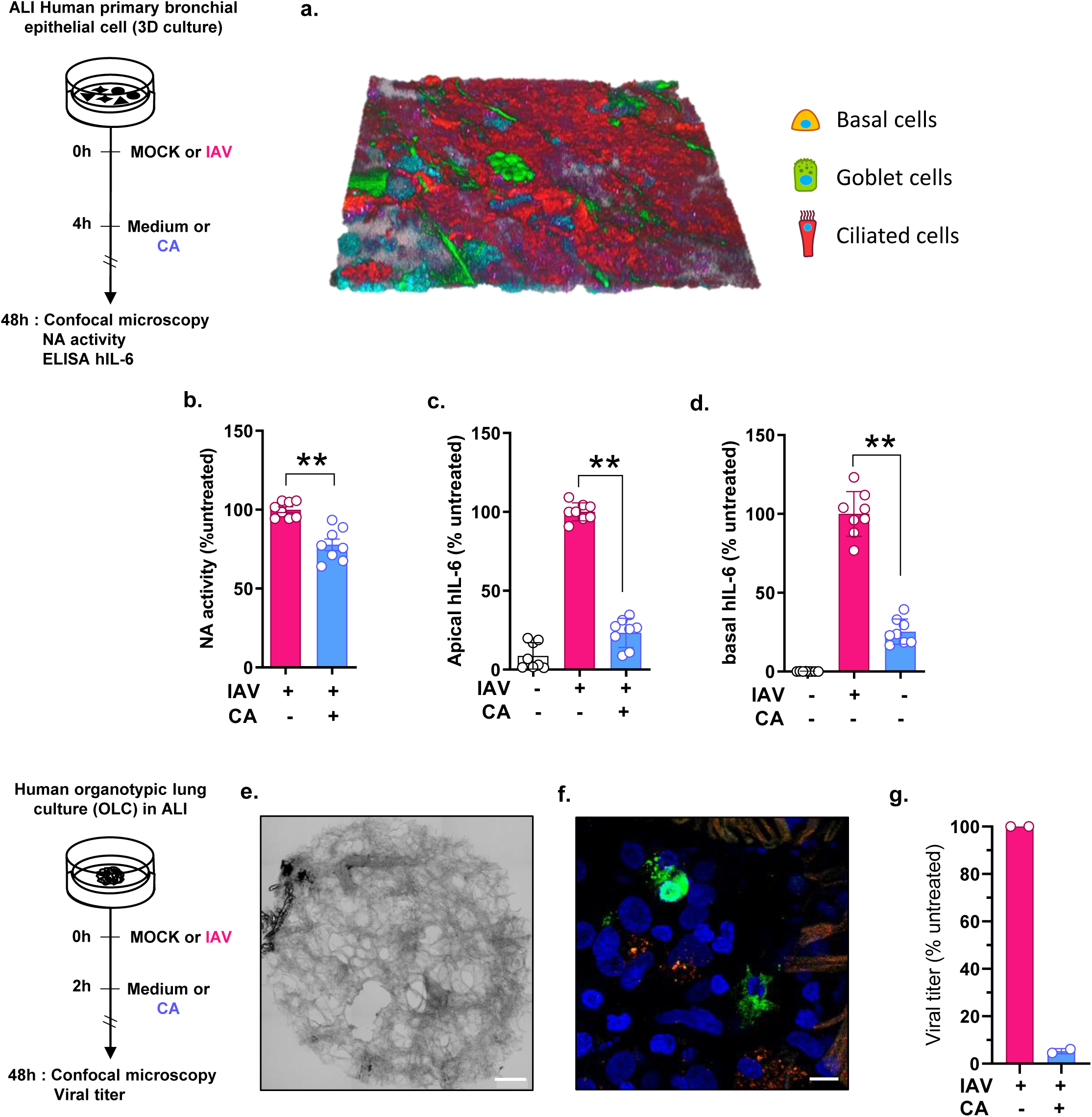
*Cis-*aco demonstrates antiviral and anti-inflammatory properties in human primary bronchial epithelial cells (PBEC) and human organotypic lung cultures (OLCs) under air-liquid interface (ALI) conditions. **(a-d)** PBEC in three-dimensional (3D) culture were infected with influenza A/Scotland/20/74 (H3N2) virus at an MOI=1 for 4 h, followed by treatment with or without 5.7 mM *cis*-aco (CA) for 44 h. **(a)** At 48 h p.i., tissue composition was validated using 3D confocal microscopy. Basal cells (yellow), goblet cells (green), ciliated cells (red) and nuclei (blue) were labeled to confirm tissue structure. **(b)** Neuraminidase activity and hIL-6 levels were measured in apical **(c)** or basal **(d)** cell supernatants. (**e-g)** OLC were infected with 2 x 10^4^ pfu of influenza A/Scotland/20/74 (H3N2) virus (IAV) and treated or not with 3.4 mM of *cis*-aco at 2 h p.i.. **(e)** Alveolar structures were visualized using light microscopy (scale bar: 500 µm). **(f)** IAV infection was confirmed *via* confocal microscopy, with viral nucleoprotein (green), nuclei (blue) and α-tubulin (red) labeled at 48h p.i. (scale bar: 10 µm). **(g)** Viral titers were measured in OLC supernatants at 48 h p.i.. Data are presented as the mean ± SEM. PBEC data **(a:** microscopy) represent duplicates from 3 independent patients analyzed in a single experiment. Panels **(b-d)** represent 4 independents experiments. Representative images of OLC from 2 patients are shown in **(e-f)** and viral titers in (**g**) were measured using pooled supernatants from 5 OLC derived from 2 patients. Statistical analyses were conducted using the Wilcoxon test **(b**) or Kruskal-Wallis test **(c-d)**.

**Figure 3.**
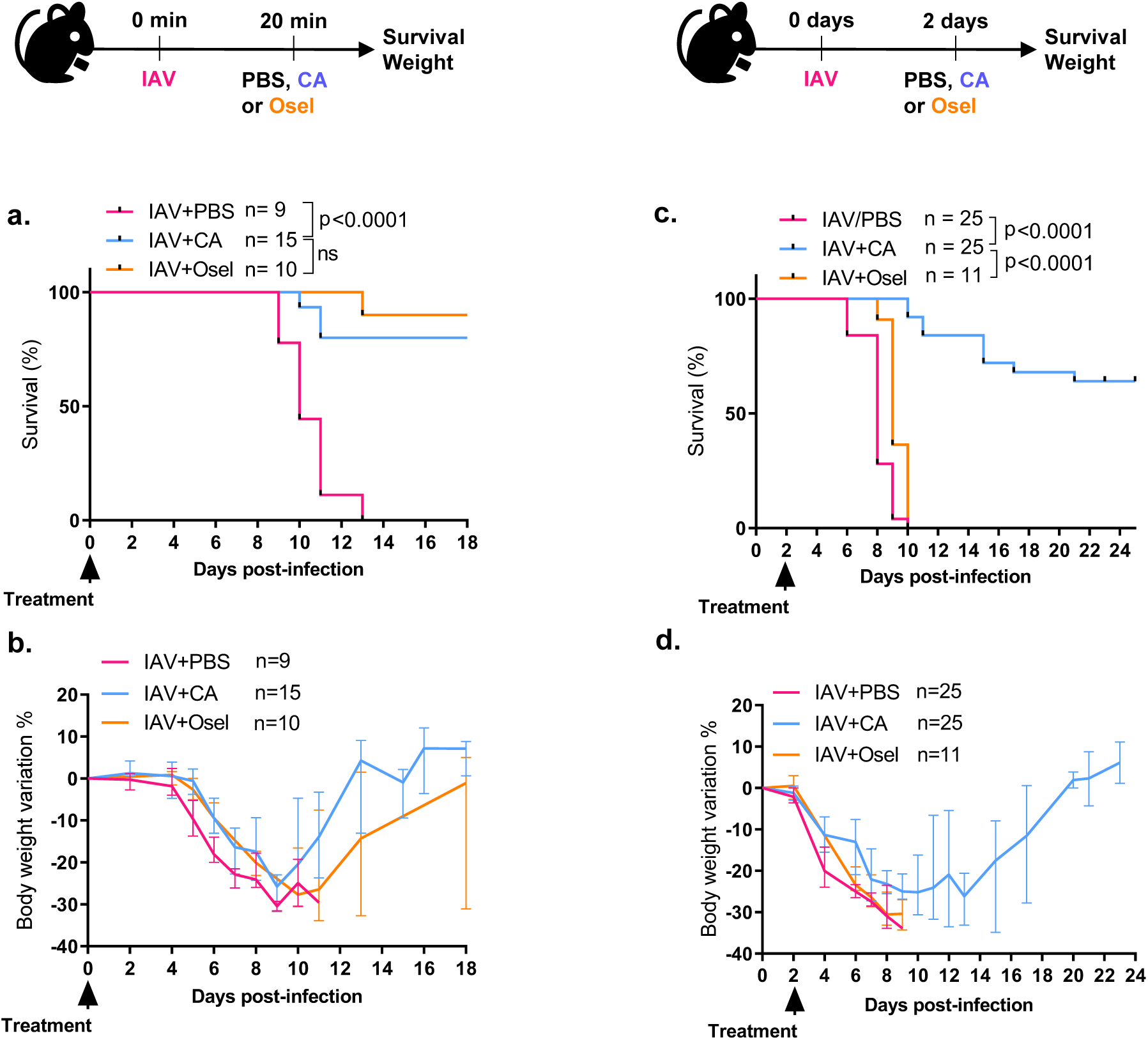
*Cis*-aco provides superior protection against influenza infection in mice compared to Oseltamivir, even with delayed treatment. Seven weeks old female C57Bl/6 mice were intranasally infected with 200 pfu of influenza A/Scotland/20/74 (H3N2) virus (IAV) and treated intranasally with either 30 mg/kg of *cis*-aco (CA, blue) or 20 mg/kg of Oseltamivir (Osel, orange) at: **(a-b)** 20 min p.i. or **(c-d)** 2 days p.i. Animal survival rates **(a,c)** and body weight loss **(b,d)** were monitored daily. Data are presented as the mean ± SEM from at least 2 independent experiments. The number of mice in each treatment group (“n”) is indicated. Statistical analysis was performed using the Log-rank (Mantel-Cox) test.

To further model human lung physiopathology, we developed a standardized *ex vivo* lung organotypic slice culture at the ALI (Fig. 2e). Tissue explants were obtained from patients undergoing surgery for lung cancer (Table 1), predominantly male (6/8), with an average age of 73 [68–78] years and a history of smoking (6/8).Upon tissue infection, IAV localized to restricted areas (Fig. 2f), and infectious particles were detected in culture supernatants (Fig. 2g). *Cis*-aco treatment reduced viral titers by 95% (Fig. 2g).

### *Cis*-aco confers potent benefits in a murine model of influenza infection

Before evaluating the potential anti-influenza effects of *cis*-aco *in vivo,* we assessed its safety profile using repeated instillations in mice over 15 days. *Cis*-aco did not significantly affect body weight in either male or female mice compared to controls (Fig. EV4a). No changes in liver function were observed, as measured by serum ALAT levels (Fig. EV4b). Similarly, no alterations in fecal microbiota composition were detected between *cis*-aco-treated and control mice (Fig. EV4c). Flow cytometry analysis of immune cells in BAL fluid (Fig. EV4d) and blood (Fig. EV4e) confirmed that *cis*-aco had no impact on pulmonary or systemic immune responses, regardless of sex.

By mitigating various aspects of influenza pathogenesis, the preceding *in vitro* and *ex vivo* data (Figs. 1 and 2) suggest that *cis*-aco could prevent lung damage induced by IAV *in vivo*. Therefore, to assess its potential as an anti-influenza drug, we evaluated its effect on the mortality of infected mice, comparing it to the standard of care, oseltamivir (a neuraminidase inhibitor [25]). While all untreated IAV-infected mice succumbed, those treated with 30 mg/kg *cis*-aco or 20 mg/kg oseltamivir within 20 min p.i. reached survival rates of 80% and 90%, respectively (Fig. 3a). Moreover, by 15 days post-treatment, surviving mice had regained their original body weight (Fig. 3b). However, delaying oseltamivir administration at 2 days p.i. abolished its efficacy (Fig. 3c-d), whereas *cis*-aco retained its curative effect under the same conditions in both female C57Bl/6 (Fig. 3c-d) and male BALB/c mice (Fig. EV5).

### *In vivo* assessment of pathophysiological mechanisms modulated by *cis*-aco in influenza pneumonia

To better understand how *cis*-aco achieves its protective effect on survival, we next explored its impact on the pathophysiological responses associated with influenza infection (Fig. 4). Mice were intranasally infected with a lethal dose of influenza A/Scotland/20/1974 (H3N2) and treated with *cis-*aco two days p.i.. Four days p.i., some mice were euthanized to analyze early events in influenza pathophysiology. *Cis*-aco treatment resulted in a significant reduction in viral load in lung tissues, with a 1-log decrease compared to controls (p<0.0001; Fig. 4a). Additionally, the relative expression of 50 mediators in the BAL fluids – including pro-inflammatory cytokines, interferons, growth factors, and proteases – was reduced in *cis*-aco-treated animals (Fig. 4b). Among these, the most pronounced reductions were observed in CCL2/MCP-1, C1qR1, G-CSF, CCL17/Tarc, and CCL20/MIP3α, while levels of resistin, myeloperoxidase, pentraxin 2/serum amyloid P, CXCL-10/IP-10, and CXCL9/Mig remained largely unchanged.

**Figure 4.**
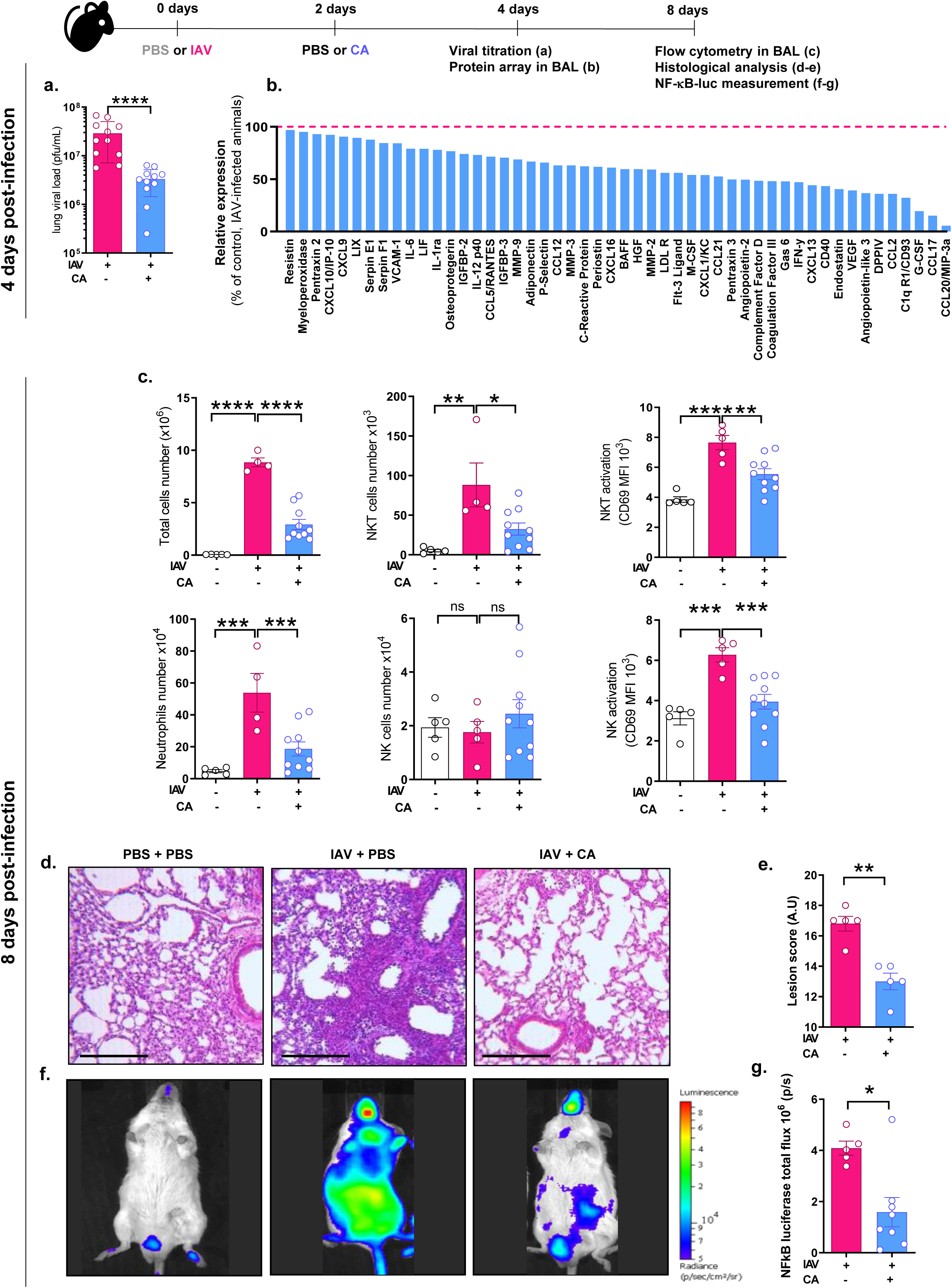
*Cis*-aco reduces viral load, lung inflammation and tissue damage in IAV-infected mice. **(a-e)** Seven-week-old female C57Bl/6 mice were intranasally infected with 200 pfu of influenza A/Scotland/20/74 (H3N2) virus (IAV) and treated or not 2 days p.i. with 30 mg/kg of *cis*-aco (CA) intranasally. Mice were euthanized at either 4 or 8 days p.i.. **(a-b)** At day 4 p.i., **(a)** the viral load in lung tissues was quantified using a PFU assay, and **(b)** the levels of 50 mediators were measured in BAL fluids. **(c-e)** At day 8 p.i., **(c)** the number and activation status of immune and inflammatory cells in BAL fluids were determined by flow cytometry. **(d-e)** Lung sections were stained with H&E and tissues lesions were assessed and scored (scale bar: 200 µm). (**f-g)** NF-κB transgenic Balb/c mice (n=9) were intranasally infected with 300 pfu of IAV and treated 2 days p.i. with 30 mg/kg CA. At 8 days p.i., mice were anesthetized, and luciferin was administered intranasally (0.75 mg/kg). Bioluminescence was quantified using the IVIS imaging system. Statistical analyses were performed using the one-way ANOVA test **(c)**, or the Mann-Whitney test **(a, e, g)**. Data are presented as the mean ± SEM and include results from 1 experiment **(b,f)** or 3 **(a)** or 2 **(c,d,e,g)** independent experiments.

In a separate series of experiments, mice were sacrificed 8 days p.i. to assess lung injury caused by IAV. Severe pathology was characterized by increased leukocytes infiltration, including neutrophils and NKT cells. While NK cell counts remained stable, their activation, assessed by CD69 marker quantification, was elevated. Similarly, NKT activation was also significantly heightened upon IAV infection (Fig. 4c). *Cis-*aco treatment consistently mitigated these responses, significantly reducing neutrophil and NKT cell infiltration (p=0.001 and p=0.05, respectively; Fig. 4c). Furthermore, NK and NKT cell activation was markedly lower in *cis*-aco-treated mice compared to infected, untreated controls (p=0.01; Fig. 4c).

Histopathological analysis revealed significant reductions in alveolar wall thickening, hyaline membrane formation, epithelial necrosis, and leukocyte infiltration in IAV-infected mice treated with *cis*-aco compared to untreated animals (Fig. 4d). Quantitative assessment confirmed that *cis*-aco significantly alleviated tissue damage caused by IAV infection (p<0.01; Fig. 4e). To further evaluate IAV-triggered inflammation on a broader scale, we measured NF-κB activity in NF-κB-luciferase transgenic mice infected with IAV and treated or not with *cis-*aco (Fig. 4f and 4g). By day 8 p.i., both airway and systemic inflammation were visible (Fig. 4f, central picture). In contrast, NF-κB activation was significantly reduced in *cis*-aco-treated animals (p<0.03; Fig. 4f, right picture, and 4g). Overall, *cis*-aco suppressed critical aspects of influenza pathogenesis, including viral replication, inflammatory signaling, secretion of inflammatory mediators, and immune cell infiltration.

### Mechanistic insights into the anti-influenza action of *cis*-aco in inhibiting influenza virus replication and inflammatory signaling

To gain molecular insights into how *cis*-aco inhibits the production of viral particles, we analyzed its effects on various stages of IAV replication cycle. IAV is an enveloped virus with a genome made up of negative sense, single-stranded RNA and its life cycle comprises the following stages: (i) entry into the host cell; (ii) nuclear import of ribonucleoproteins (vRNP); (iii) transcription of the viral genome; (iv) replication and translation of viral proteins; (v) nuclear export of vRNP; and (vi) assembly and budding at the host cell plasma membrane.

#### Impact of cis-aco on virus budding and viral protein expression

Transmission and scanning electron microscopy (TEM and SEM) of IAV-infected bronchial epithelial cells revealed a marked reduction in virus budding following *cis*-aco treatment (Fig. 5a). To determine whether this was due to impaired budding or to reduced viral material production, we quantified viral proteins and mRNA in cells, treated or not with *cis*-aco 4 h p.i.. Confocal microscopy (Fig. 5b-c) and western-blotting (Fig. 5d-e) revealed significant reductions (∼75%, *p*<0.03, Fig. 5e) in NP, NS1, and PA protein expression at 8 h p.i. while qRT-PCR showed a 10-fold decrease in viral M1 mRNA levels at 6 h p.i. (p<0.02, Fig. 5f). Given that *cis*-aco was applied at 4 h p.i., it is unlikely that it affects the early stages of viral entry, including cell fusion and nuclear trafficking (completed within 1 h p.i., [26]). These findings suggest that *cis*-aco inhibits IAV genome transcription, leading to reduced viral protein expression and neo-virion production. This hypothesis was confirmed by a minigenome assay in HEK293T cells, where *cis*-aco treatment inhibited luciferase activity by ∼75% (p < 0.03, Fig. 5g), indicating interference with IAV polymerase activity.

**Figure 5.**
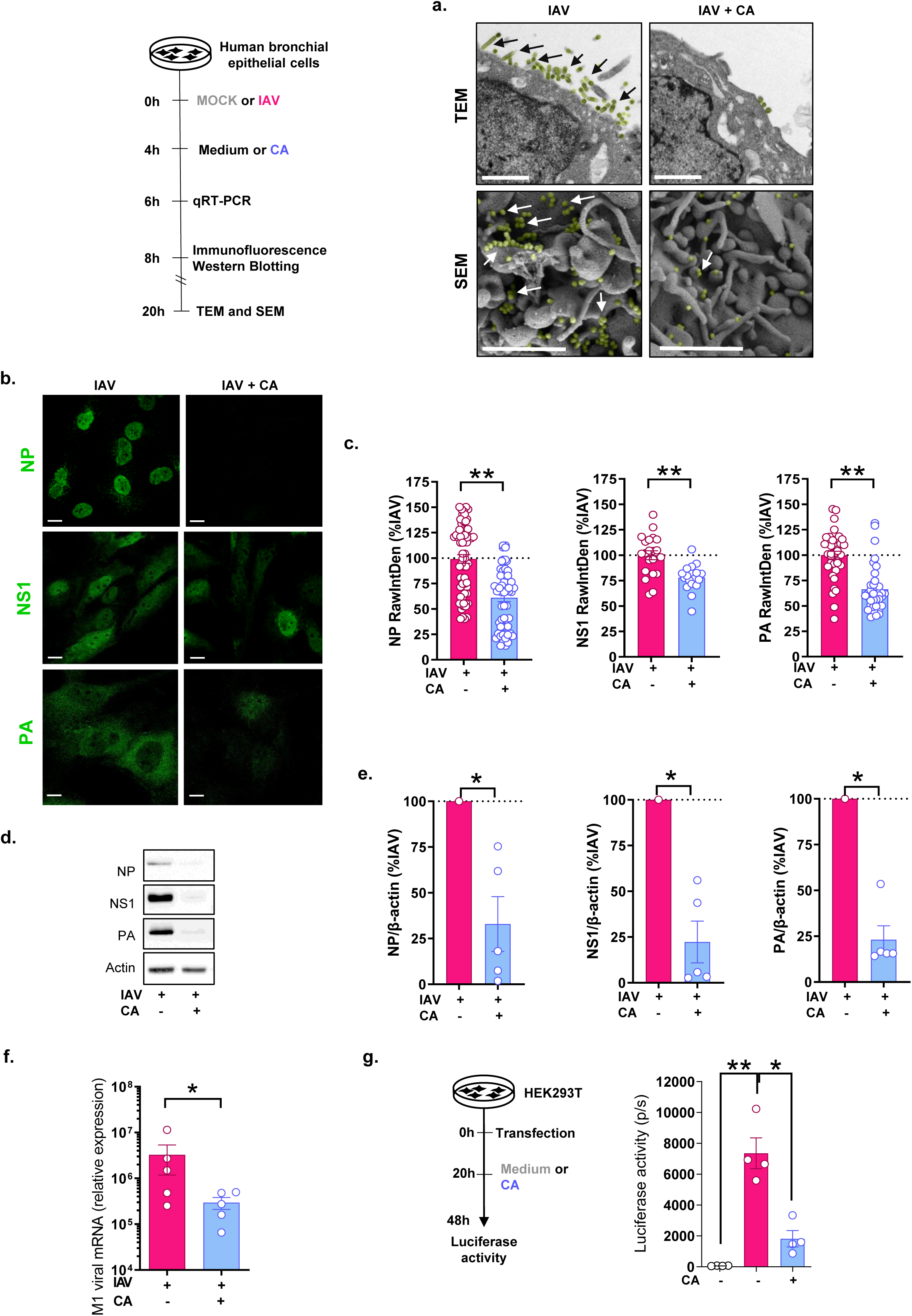
Anti-influenza virus properties of *cis*-aco involve the inhibition of viral polymerase activity. BEAS-2B cells were infected with influenza A/Scotland/20/74 (H3N2) virus at a MOI=5 **(a)** or MOI=1 **(b-f)** for 4 h, then washed and treated with 3.4 mM of *cis*-aco (CA) or left untreated (Medium). (**a)**: Representative images from transmission electron microscopy (upper panel) and scanning electron microscopy (lower panel) show IAV particles budding at 20 h p.i., indicated by arrows (scale bar: 1 µm). **(b-e)** At 8 h p.i., viral protein (green) expression and trafficking were analyzed by **(b-c)** confocal microscopy (scale bar: 20 µm) and **(d-e)** Western blotting to detect viral NP, NS1, and PA proteins. **(c)** Raw integrated density (RawIntDen), calculated as the sum of all pixel values in the region of interest, was measured and normalized to the mean of the IAV condition for each experiment. **(e)** Relative protein levels were normalized to the mean value of “IAV condition” samples, with β-actin as a loading control. **(f**) At 6 h p.i., IAV transcription was quantified by RT-qPCR, measuring M1 viral mRNA levels. **(g)** A minigenome assay was performed in HEK-293T cells to test the effect of *cis*-aco on viral polymerase activity. Cells were transfected with plasmids encoding PA, PB1, PB2, NP, and the reporter plasmid pPolI-WSN-NA-firefly luciferase. At 20 h post–transfection, cells were treated with 0 or 3.4 mM *cis*-aco (CA) and luciferase activity was measured at 48h post-transfection. Results are presented as the mean ± SEM from 3 **(a, b, c)**, 4 **(d, e, g),** or 5 **(f)** independent experiments. Statistical analyses were performed using the Kruskal-Wallis test with Dunn’s multiple comparison test **(g)**, the Mann-Whitney test **(c)**, or the Wilcoxon matched-pairs rank test **(e,f)**.

#### Anti-inflammatory effects of cis-aco

We next investigated whether *cis*-aco’s ability to decrease inflammation (e.g., IL-6, Figs. 1c-d) was a secondary consequence of its antiviral action or due to intrinsic immunomodulatory properties.

IAV infection in lung epithelial cells activates intracellular signalling pathways that upregulate the expression of inflammatory cytokines [27–29]. Consistent with this, we observed increased phosphorylation of extracellular signal-regulated kinase (ERK)1/2, Protein kinase B (PKB; also known as AKT), and the p65 subunit of the nuclear factor-kappa B (NF-κB) at 20 h p.i compared to non-infected cells (MOCK) (Fig. 6a). *Cis-*aco treatment appeared to inhibit the accumulation of these phosphorylated proteins (Fig. 6a-b). To further assess whether *cis*-aco possesses inherent anti-inflammatory properties, we stimulated lung epithelial cells with different inflammatory agonists: (i) Poly(I:C) (PIC), a TLR3 agonist that mimics viral RNA (ii) Phorbol 12-myristate 13-acetate (PMA), which activates protein kinase C signaling; and (iii) TNFα, a major inflammatory cytokine. As expected, all agonists induced IL-6 secretion (Fig. 6c). Interestingly, *cis*-aco treatment inhibited IL-6 release in a dose-dependent manner (2.3-fold decrease after TNFα stimulation, p<0.04, and >4-fold decrease after PIC or PMA stimulation, *p*<0.01; Fig. 6c). These findings highlight the intrinsic anti-inflammatory properties of *cis*-aco, independently of its antiviral activity.

**Figure 6.**
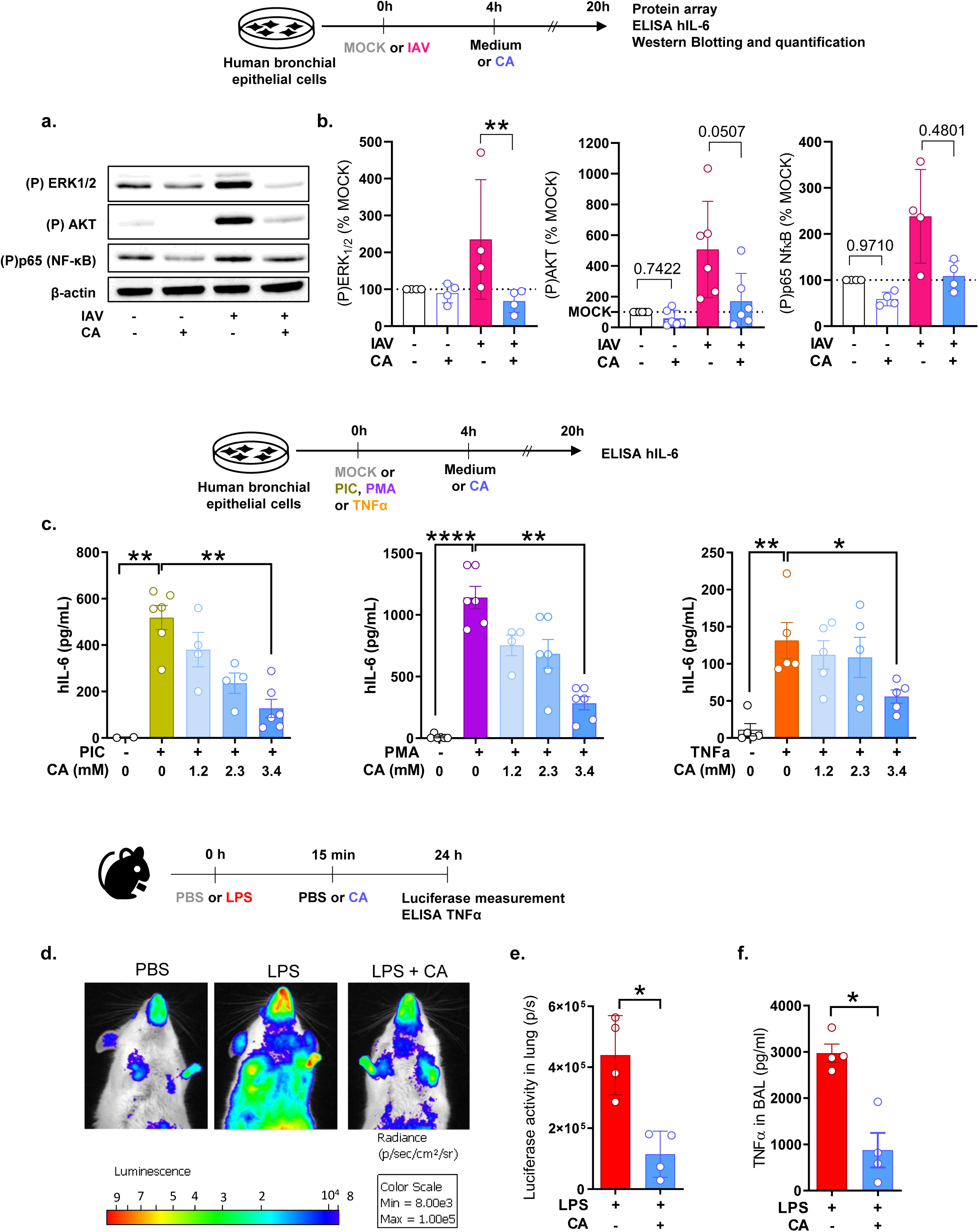
*Cis*-aco reduces pro-inflammatory responses and signaling. BEAS-2B cells were infected or not with the influenza A/Scotland/20/74 (H3N2) virus at an MOI=1 for 4 h, and subsequently treated or not with 3.4 mM of *cis*-aco (CA) for 16 h **(a-b)**. **(a)** Representative Western-blotting showing phosphorylated forms (P) of ERK1/2, AKT and p65 proteins with β-actin as a loading control. **(b)** Signal quantification of proteins and normalization was performed for each experiment. **(c)** BEAS-2B cells were stimulated or not (MOCK) with 2 µg/mL Poly(I:C) (PIC), 50 nM Phorbol 12-myristate 13-acetate (PMA), or 20 ng/mL Tumor necrosis factor alpha (TNFα) for 4 h, followed by treatment with increasing doses of *cis*-aco (CA) for 16 h. IL-6 levels in cell supernatants were measured by ELISA. **(d-f)** NF-κB transgenic Balb/c mice were instilled with 10µg LPS and treated intranasally with either PBS or 30 mg/kg *cis*-aco (CA) 15 min post-stimulation. At 24h post-instillation, bioluminescence was measured using the IVIS imaging system after intranasal administration of luciferin. Data are presented as the mean ± SEM. Results reflect cumulative data from a single experiment (**d-f;** n=4 mice *per* condition) or at least 4 independent experiments (**a-c)**. Statistical analyses were performed using the Friedman test **(b)**, Kruskal-Wallis test (**c**), or Mann-Whitney test **(e-f)**

#### In vivo confirmation of the anti-inflammatory activity of cis-aco

To examine the anti-inflammatory effects of *cis*-aco *in vivo*, we used a murine model of acute lung injury induced by bacterial lipopolysaccharide (LPS; [30]). To this end, NF-kB-luciferase transgenic mice were treated or not with *cis*-aco 15 min post-LPS challenge, and NF-kB activity was measured as a proxy of inflammation (Fig. 6d-e). At 24 h post-stimulation, lung inflammation induced by LPS (Fig. 6d, center picture) was significantly reduced in *cis*-aco-treated animals (∼75%, *p*<0.03; Fig. 6d, right picture and Fig. 6e). Consistent with this observation, TNFα levels in the BAL fluids of LPS-challenged mice were also significantly lower in *cis*-aco-treated animals (70% reduction, *p*<0.008, Fig. 6f). These *in vivo* experiments thus suggest that *cis*-aco can inhibit inflammatory signaling, resulting in decreased production of pro-inflammatory cytokines.

### The anti-influenza properties of *cis*-aco are independent of itaconate

*Cis*-aco is a TCA intermediate that can be converted into itaconate by the mitochondrial enzyme *cis*-aco decarboxylase (CAD), also known as ACOD1 or Irg1 (Fig. 7a) [31]. To verify whether the anti-influenza effect of *cis*-aco is dependent on this conversion, we used a silencing approach targeting the CAD gene. This specific siRNA transfection effectively suppressed CAD expression in bronchial epithelial BEAS-2B cells compared to a control scramble siRNA (Fig. 7b. Interestingly, even under CAD silencing conditions, *cis*-aco retained its antiviral and anti-inflammatory properties, as determined through quantification of IAV infectious particles, neuraminidase activity, and IL-6 release (Fig. 7c-e). These findings suggest that *cis*-aco inhibits IAV infection in lung epithelial cells independently of its conversion into itaconate by CAD. These *in vitro* findings were further supported by *in vivo* experiments, where CAD-deficient mice infected intranasally with IAV and treated with *cis*-aco displayed survival rates similar to wild-type mice (Fig. 7f) [32].

**Figure 7.**
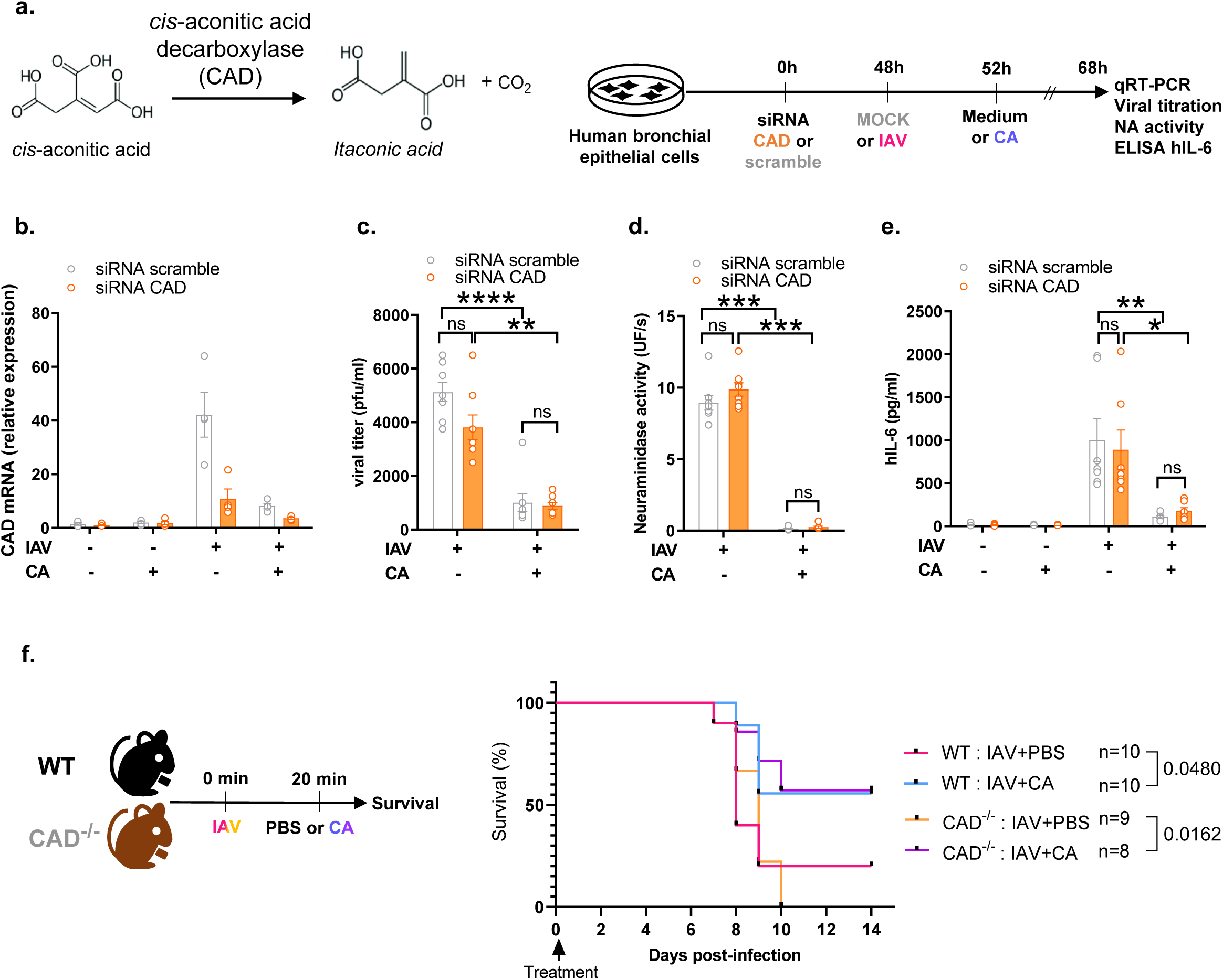
Anti-influenza activity of *cis*-aco is independent from itaconate. **(a-e)** BEAS-2B cells were transfected with either *cis*-aconitate decarboxylase (CAD) or control (scramble) siRNA. At 48 h post-transfection, cells were infected or not with A/Scotland/20/74 (H3N2) virus (IAV) at an MOI=1 for 4 h, and subsequently treated or not with 3.4 mM *cis*-aco (CA) for 16h. **(a)** CAD catalyzes the decarboxylation of *cis*-aco to produce itaconate. **(b)** Gene knockdown was confirmed by RT-qPCR. **(c, d)** IAV particles production was measured by a plaque-forming units (pfu) assay **(c)** and a neuraminidase activity assay **(d)**. **(e)** IL-6 levels in cell supernatants were quantified by ELISA. **(f)** CAD-deficient and wild-type (WT) mice were infected intranasally with 100 pfu of A/Scotland/20/74 (H3N2) virus (IAV) and treated intranasally or not with 30 mg/kg of *cis*-aco (CA) 20 min p.i.. Animal survival was monitored daily. Data are presented as the mean ± SEM and are cumulative from a single experiment **(f)**, or 3 **(b),** or 4 **(c-e)** independent experiments. Statistical analyses were performed using the Kruskal-Wallis test with Dunn’s multiple comparison test **(a-e)**, or the Log-rank (Mantel-Cox) test **(f)**.

### *Cis*-aco mitigates mortality in IAV-infected mice in a clinically relevant timeframe

The preceding data demonstrate that *cis*-aco prevents influenza-induced lung damage by reducing viral infection and inflammation. To assess its therapeutic potential in a clinically relevant context, we considered the typical delay between symptom onset and treatment in human influenza cases [33]. This delay is critical for evaluating the efficacy of anti-influenza therapies in mice in “real-world” conditions. To complement our experimental study, we examined a prospective clinical trial [34] involving patients with community-acquired pneumonia (CAP) caused by influenza A and B viruses. Among the 153 CAP patients, 37% had viral pneumonia, 24% had bacterial pneumonia and 20% had co-infections. IAV was the predominant pathogen, found in 33% of cases. Detailed characteristics of these CAP patients, particularly those with IAV infection, are provided in Table 2. The median [IQR] symptom-to-hospitalization time was 3 [2–7] days for all CAP cases and 4 [3–6] days for those attributed to influenza (A or B) infection (Fig. 8a).

**Figure 8.**
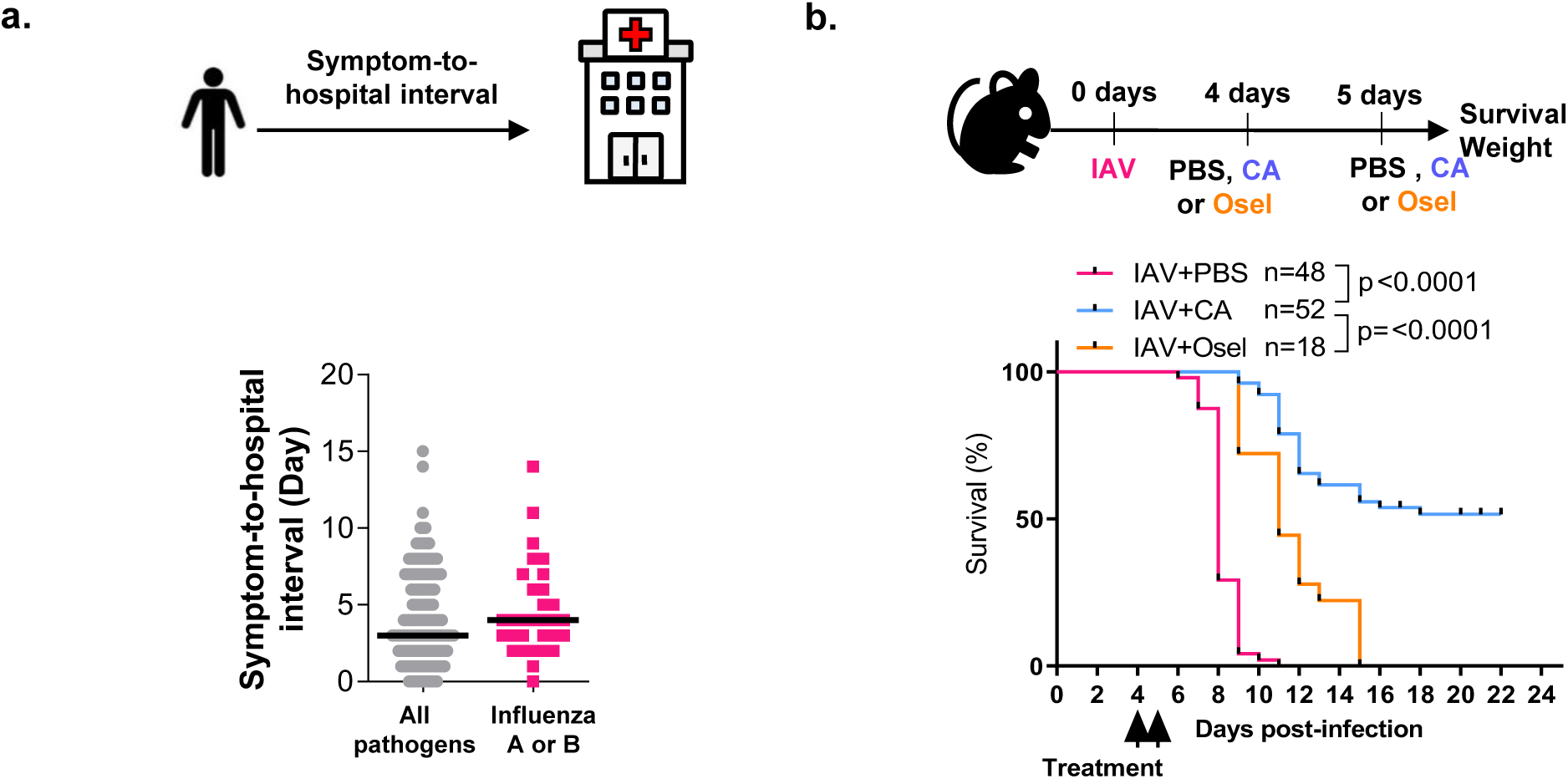
*Cis*-aco protects mice from influenza infection within a clinically relevant time frame. **(a)** Time between the symptoms onset and the first hospital admission in patients hospitalized for community acquired pneumonia (CAP). Each dot represents an individual patient with the the median indicated by the black line. **(b)** Seven-week-old female mice were intranasally infected with 200 pfu of influenza A/Scotland/20/74 (H3N2) virus (IAV) and treated intranasally 4 and 5 days p.i. with 30 mg/kg *cis*-aco (CA, blue) or 20mg/kg Oseltamivir (Osel, orange), or left untreated (pink). Survival was monitored daily. Data in **(b)** are presented as the mean ± SEM and are cumulative from 3 independent experiments; the number of mice (“n”) is indicated. Statistical analysis was performed using the Log-rank (Mantel-Cox) test.

To simulate this clinically relevant delay, *cis*-aco treatment was initiated in mice on day 4 p.i., by which time IAV infection had already progressed to pneumonia, with substantial production of inflammatory mediators (Fig. 4b). To more closely mirror clinical conditions, a second dose of *cis*-aco was administered on day 5 p.i.. In line with previous clinical observations [35, 36] and reinforcing our data from Fig. 3, delayed oseltamivir treatment did not prevent IAV-induced mortality (Fig. 8b). In contrast, *cis*-aco treatment on days 4 and 5 improved survival, increasing the survival rate from 0% to approximately 50% (Fig. 8b). These findings highlight the potent curative effects of *cis*-aco against IAV infection within a clinically relevant timeframe.

## DISCUSSION

Influenza pathophysiology is multifactorial, involving direct viral cytopathic effects and dysregulation of the host immune response, which together contribute to lung damage. Currently, the treatment of influenza remains an unmet medical need, as the effectiveness of antiviral therapies is limited, especially when administered late in the course of infection. Moreover, strategies that reduce inflammation without controlling viral replication have been associated with increased mortality [37]. In this study, we show that *cis*-aco treatment addresses both key aspects of influenza pathophysiology. *Cis*-aco effectively inhibits major human influenza strains by impairing viral RNA and protein expression, while simultaneously reducing influenza-induced inflammatory signaling. Guided by a clinical perspective, we demonstrated the protective anti-influenza effects of *cis*-aco in human lung tissue explants and in infected mice treated within a clinically relevant timeframe, surpassing the efficacy window of the standard of care, oseltamivir/Tamiflu.

By screening a range of related metabolites using an *in vitro* model of IAV infection in human bronchial epithelial cells, the primary target cells for IAV [38–40], *cis*-aco emerged as the most promising molecule. It is synthesized in the TCA cycle from citric acid through the action of the mitochondrial enzyme aconitate hydratase (also named aconitase) and can be converted by the enzyme *cis*-aconitate decarboxylase (CAD; also called ACOD1 or Irg1) into itaconate. This latter metabolite is central in linking the innate immune response to cell metabolism, including during IAV infection [31, 41, 42]. However, the anti-influenza effects of *cis*-aco were not due to its conversion into itaconate, as antiviral and anti-inflammatory activities persisted despite efficient silencing of CAD using siRNA. These *in vitro* findings were further confirmed *in vivo*, with CAD-deficient mice infected intranasally with IAV and treated with *cis*-aco displaying survival rates comparable to wild-type mice. Collectively, these data demonstrate that the anti-influenza properties of *cis*-aco are independent of its conversion to itaconate.

To further elucidate the mechanism of action of *cis*-aco, we investigated its effects on various stages of the IAV life cycle. Our results show that *cis*-aco impairs viral polymerase activity, leading to decreased levels of viral RNA and proteins and preventing the formation of new viral particles. These antiviral effects were confirmed against both influenza A and B viruses, which is particularly noteworthy given that current neuraminidase inhibitors are less effective against influenza B viruses compared to influenza A [5, 43]. However, further research is required to fully elucidate the mechanism of *cis*-aco, given its likely complex effects on both viral replication and inflammatory cell signaling.

Severe viral pneumonia is closely associated with lung hyperinflammation, which can lead to acute respiratory distress syndrome (ARDS), as well as significant morbidity and mortality due to respiratory failure. While antiviral therapy and supportive measures, such as mechanical ventilation, are standard treatments, the use of immune-modulatory agents remains controversial [37]. Proponents argue that anti-inflammatory treatments could reduce lung inflammation whereas opponents caution that such treatments might interfere with immune responses, potentially delaying viral clearance and increasing mortality risk. This debate was particularly evident during the A/H1N1 2009–2010 influenza pandemic, when anti-inflammatory therapies like steroids were linked to higher mortality rates [44–46]. As a result, anti-inflammatory therapies are not recommended for influenza-related pneumonia, and the disease was excluded from the largest randomized clinical trial assessing steroids in severe community-acquired pneumonia (*i.e*. the CAPE COD study)[47].

Given this context, *cis*-aco’s dual action as both an antiviral and anti-inflammatory agent holds considerable therapeutic potential. By simultaneously reducing hyperinflammation and enhancing viral clearance, *cis*-aco could offer a comprehensive treatment strategy. It exerts its anti-inflammatory effects by inhibiting key pro-inflammatory pathways, including ERK, AKT, and NF-κB, which are also crucial for viral replication and often hijacked by IAV [48–55]. For instance, IAV activation of the ERK pathway facilitates the nuclear export of vRNPs [51–54]. By modulating these cellular signaling factors, *cis*-aco may impair IAV replication more potently than existing antiviral drugs that primarily target viral components [56–58]. Furthermore, because *cis*-aco acts on cellular pathways rather than solely targeting viral factors, it has the potential to limit the risk of drug resistance by reducing selective pressure on the virus itself [59].

To confirm the protective mechanisms of *cis*-aco in a more representative and challenging context, we conducted *in vivo* experiments using mice infected with a lethal dose of IAV [16, 60]. *Cis*-aco mitigated all key aspects of influenza pathology, including reducing viral replication, controlling excessive inflammatory cytokine production, decreasing immune cells recruitment and activation, and minimizing tissue lesions.

Beyond its efficacy against infectious diseases, our study also emphasizes the broader therapeutic potential of *cis*-aco. Its potent anti-inflammatory properties position it as a promising candidate for managing non-infectious pulmonary inflammatory conditions, such as asthma or fibrosis. Local administration of *cis*-aco locally *via* inhalation could enhance its therapeutic effects while reducing potential systemic side effects [66].

Preclinical studies are essential for advancing potential treatments but traditional models – such as mouse and other animal models or *in vitro* cell cultures – often fail to accurately predict efficacy in patients. To improve clinical translation, we used a multimodal strategy that included: (*i*) human lung samples from individuals at high risk for influenza (*e.g*. elderly), (*ii*) a dosing schedule mimicking clinical treatment (instead of prophylactic or simultaneous treatment, as typically used), and (*iii*) comparison with an FDA-approved drug. We demonstrated *cis*-aco antiviral effects not only in human bronchial epithelial cell lines but also in advanced *ex vivo* lung culture models such as primary epithelial cells and human organotypic lung cultures. The latter preserves the complex lung tissue architecture and cellular diversity, providing a more relevant evaluation context than conventional cell cultures [61]. Next, we compared *cis*-aco with oseltamivir, the most commonly recommended anti-influenza drug [20]. While oseltamivir is effective at early infection stages, its limited therapeutic window [62, 63] and delayed administration in many patients restricts its efficacy. For example, only 20% of patients receive oseltamivir within 2 days of symptom onset, while viral pneumonia hospitalization typically occurs 4-5 days post-symptom onset [36, 64]). These findings highlight the importance of evaluating anti-influenza treatments at day 4 post-infection to better reflect clinical realities.

Interestingly, in our IAV-infected mouse model, oseltamivir’s therapeutic effect declines markedly when administered at 2 or 4 days p.i., aligning with observations in critically ill influenza patients [36]. In contrast, *cis*-aco demonstrated a significant survival benefit, with ∼70% survival when administered at day 2 p.i. and ∼50% survival at day 4. These results are promising as similar efficacy is rarely achieved with FDA-approved influenza drugs under similar experimental conditions [65]. The superior efficacy of *cis*-aco across a larger treatment window underscores its potential as a major advancement in influenza therapy, allowing greater flexibility and effectiveness.

Our current findings significantly advance the identification and understanding of host-derived metabolites with antiviral activity, building on our previous studies on succinate[16]. Succinate mediates its anti-influenza effects by inducing the succinylation of a single amino acid (K87) on the IAV nucleoprotein, thereby impairing the trafficking of viral ribonucleoprotein complexes and disrupting the replication cycle. However, succinate has several limitations compared to *cis*-aco: it requires concentrations five times higher to achieve optimal anti-IAV inhibition, it is ineffective against influenza B viruses and it does not modulate the inflammatory signaling pathways associated with infection [16]. In contrast, *cis*-aco offers the advantage of dual antiviral and anti-inflammatory actions, positioning it as a more potent endogenous molecule for controlling influenza infection. Thus, *cis*-aco, succinate and other metabolites like fumarate [67] and itaconate [68] are increasingly recognized as endogenous mediators with remarkable antimicrobial activities. This discovery points to an ancient, conserved mechanism among primitive organisms, likely evolved to balance cellular proliferation and defense through a unique molecular system. This hypothesis aligns with theories on the origins of life, where metabolites were not only crucial for growth and development but also served as protective agents against environmental challenges [69]. In a broader context, our findings further support the concept that metabokines are part of a wider family of host defence mediators – including cytokines, chemokines, bioactive lipids, and reactive oxygen species – that collectively orchestrate responses to external threats [70, 71].

In conclusion, our study demonstrates that *cis*-aco, a host-derived metabolite, exhibits potent anti-influenza properties with significant translational potential due to its natural origin and anticipated low toxicity. As illustrated in Fig. 9, *cis-*aco shows: (i) antiviral activity through inhibition of the IAV polymerase, (ii) strong anti-inflammatory and anti-cell death effects, (iii) broad-spectrum action against both influenza A and B viruses, and (iv) protective efficacy surpassing that of the reference drug oseltamivir. Altogether, our results pave the way for the development of *cis*-aco-based therapies for influenza virus infections Further studies are warranted to investigate *cis*-aco pharmacokinetics and explore its potential in combination with existing antivirals.

**Figure 9.**
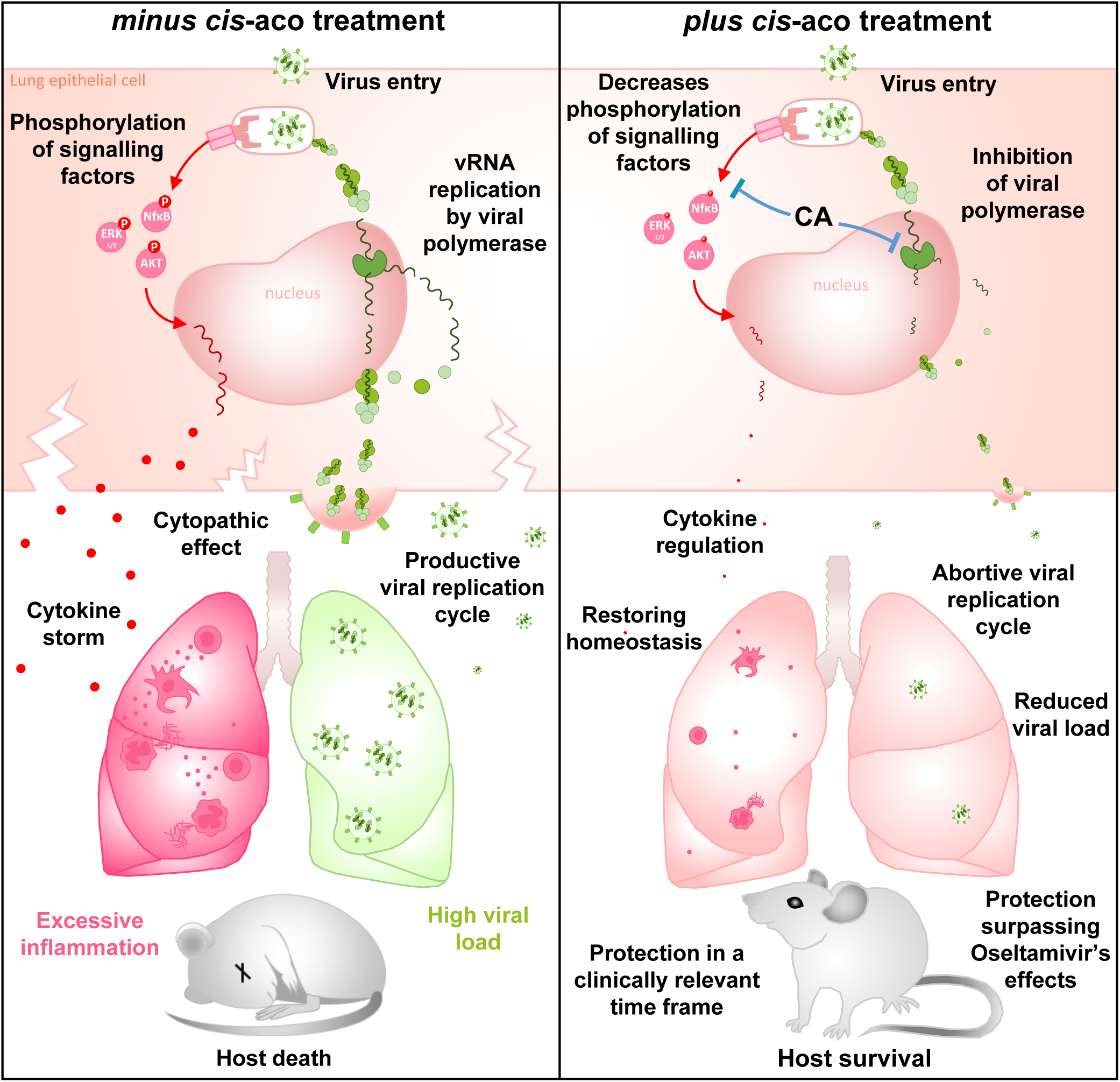
AntiD influenza mechanism of action of *cis*-aco. *Cis*-aco (CA) impairs IAV polymerase activity, reducing viral mRNA expression and protein synthesis, thereby preventing effective virus replication. Additionally, *cis*-aco downregulates inflammatory pathways triggered by various stimuli. *In vivo*, these combined antiviral and anti-inflammatory effects decrease viral load and mitigate excessive inflammation, providing superior protection against mortality compared to the reference anti-influenza drug Oseltamivir.

## METHODS

### Viruses

The influenza strains used in this study were initially provided by partner laboratories and subsequently amplified in M. Si-Tahar’s laboratory. Specifically, the mouse-adapted influenza A/Scotland/20/74 (H3N2) strain was kindly provided by Prof. Sylvie van der Werf’s team at the Pasteur Institute, Paris, France; the influenza A/PR/8/34 (H1N1) strain by Dr. Georg Kochs at Freiburg University, Germany; the pandemic H1N1 strain by Dr. François Trottein at the Center for Infection and Immunity of Lille; and the influenza B/Paris/234/2013 (Yamagata lineage) strain was obtained through the European Virus Archive Global (EVAg)

### Cell lines culture

*In vitro* experiments were performed using human bronchial epithelial BEAS-2B cells, except for plaque assays which used Madin-Darby Canine Kidney (MDCK) cells, and minigenome assay which used HEK-293T. These cells were cultured in either F-12K Medium (BEAS-2B) or MEM (HEK-293T and MDCK) supplemented with 10% FBS, 100 U/ml penicillin, and 100 µg/ml streptomycin. All cells were mycoplasma-free. BEAS-2B cells were infected in medium without FBS for 4 h with IAV at MOI=1 (except for TEM and SEM analysis, for which an MOI=5 was applied). Cells were also stimulated in medium with FBS with 2 µg/ml Poly(I:C) or in medium without FBS with 2 µg/ml PMA or with 20ng/ml TNFα. Four hours after the challenge, cells were washed with PBS and incubated for 4 h or 16 h with different concentrations of metabolites (*cis*-aconitate derivative *cis*-aconitic acid, *trans*-aconitic acid, itaconic acid, glucose, pyruvate, oxaloacetic acid, fumarate, isocitric acid) diluted in medium without FBS.

### Studies involving human participants

Informed consent was obtained from all patients in accordance with the Helsinki Declaration. Tissue and cell collections were declared to the French Ministry of Graduate Study, Research, and Innovation (DC-2008-308, MESRI). Lung lobes were collected immediately following surgical resection at CHRU of Tours. All experiments adhered to The Code of Ethics of the World Medical Association and were approved by the Ethics Committee of the CHRU of Tours. Lung donors for this study varied in age, gender, medical history, and the cause of resection (see Table 1). Prospective data collection presented in Table 2 was conducted in a single center over an 18-month period. The study complied with French law for observational studies and to the STROBE guidelines for observational studies. The study was approved by the ethics committee of the French Intensive Care Society (CE SRLF 13–28), was approved by the “Commission Nationale de l’Informatique et des Libertés” (CNIL) for the treatment of personal health data. We gave written and oral information to patients or next-of-kin. Patients or next-of-kin gave verbal informed consent, as approved by the ethic committee. Eligible patients were adults hospitalized in ICU for CAP. Pneumonia was defined as the presence of an infiltrate on a chest radiograph and one or more of the following symptoms: fever (temperatureL≥L38.0L°C) or hypothermia (temperatureL<L35.0L°C), cough with or without sputum production, or dyspnea or altered breath sounds on auscultation. Community-acquired infection was defined as infection occurring within 48 h of admission. Cases of pneumonia due to inhalation or infection with pneumocystis, pregnant women and patients under guardianship were not included. Cases with PaO2L≥L60LmmHg in ambient air or with the need for oxygen therapy ≤4LL/min or without mechanical ventilation (invasive or non-invasive) were not included.

### Human primary bronchial epithelial cells (PBEC) culture

PBEC were isolated from normal bronchial tissues of lung cancer patients undergoing lobectomy at the CHRU of Tours. The cancer-free tissues were washed and incubated for 2 h at 37°C with 0.018% (w/v) proteinase XIV in Ca2+/Mg2+-free Hank’s Balanced Salt Solution. Epithelial cells were scraped from the luminal surface, washed, and cultured in serum-free keratinocyte medium supplemented with 2.4 ng/ml epidermal growth factor, 25 µg/ml bovine pituitary extract, 1 µM isoproterenol, 100 U/ml penicillin, and 100 µg/ml streptomycin on 6-well plates coated with 30 µg/ml PureCol, 10 µg/ml bovine serum albumin, and 5 µg/ml fibronectin. During the first week, 1/500 Primocin was added to the medium. After reaching near-confluence, cells were trypsinized and stored in liquid nitrogen.

For mucociliary differentiation, PBEC were cultured in PneumaCult EX medium, with a 3-4 day proliferation step before stimulation with 2 μg/ml Poly(I:C) or 50 nM PMA. The stimulation medium was a 1:1 mixture of BEGM and complete DMEM/F12, supplemented with 100 U/ml penicillin, 100 µg/ml streptomycin, 12.5 ml 1M HEPES, and 5 ml GlutaMAX™.

For ALI differentiation, PBEC were cultured submerged on semipermeable transwell inserts (0.4 µm pore size) coated with collagen and fibronectin. Once confluent, the apical medium was removed, and cells were cultured at ALI for 3 weeks.

### Human organotypic lung culture (OLC)

The preparation of OLC was carried out according to a protocol derived from [73], using human lung resections from surgical patients at the CHRU of Tours, collected in Hibernate medium containing 1/500 Primocin. Lung explants were sliced using the McIlwain® tissue chopper (Campden Instruments) at 500 μm thickness and placed back in Hibernate medium with 1/500 Primocin for slice dissociation under a dissection microscope before their immediate transfer at ALI. Individualized OLC were placed on semipermeable Millicell® cell culture inserts with PTFE membranes (0.4 µm pore size; Merck) already pre-activated with 1 ml of OLC culture medium. The culture medium is a volume-to-volume mix of BEGM and DMEM medium with 100 U/ml penicillin and 100 µg/ml streptomycin. OLC were infected by drop deposition with 2.10^4^ pfu of A/Scotland/20/74 (H3N2) virus (IAV). After 2 h, OLC were treated with 3.4 mM of *cis*-aco. At 48h p.i., the OLC subnatants were collected to measure viral titers.

### Animal care, handling and study approvals

C57Bl/6 female mice (∼8 weeks old) were purchased from Centre d’Elevage R. Janvier (Le Genest Saint-Isle, France) and housed under specific-pathogen-free conditions at Tours University animal facility, (France), with *ad libitum* access to food and water. NF-κB luciferase transgenic BALB/C mice were generated by backcrossing NF-κB luciferase transgenic B10.A mice (a kind gift from Prof. Richard Flavell, Howard Hughes Medical Institute) with BALB/C mice to produce transgenic mice with white fur, minimizing light absorption. C57BL/6NJ wild-type mice and C57BL/6NJ-Acod1em1(IMPC)J/J (CAD-/-) mice, deficient in CAD expression, were purchased from The Jackson Laboratory (Bar Harbor, ME, USA). All mice were maintained and bred at the Pasteur Institute of Lille, France (agreement B59-350009).

All procedures involving C57Bl/6 mice were conducted in compliance with European animal welfare regulations. Experiments adhered to the ethical standards set by the French government and were approved by our local and national ethics committees (CEEA.19, APAFIS#201604071220401.V2-4885, 2016111512369894 V3 – 7590). Studies using NF-κB transgenic BALB/c mice were approved by the Animal Care and Use Committee at the “Centre de Recherche de Jouy-en-Josas” (COMETHEA) under the relevant institutional authorization (Ministère de l’éducation nationale, de l’enseignement supérieur et de la recherche), authorization number: 2015100910396112v1 (APAFIS#1487). Experiments involving C57BL/6NJ-Acod1em1(IMPC)J/J (CAD-/-) mice deficient in CAD expression were ethically approved by the French Committee on Animal Experimentation and the Ministry of Education and Research (APAFIS#10232-2017061411305485 v6, approved on 14/09/2018).

### Neuraminidase (NA) assay

The assay measures the release of a 4-methylumbelliferone fluorescent product from the 2′-(4-Methylumbelliferyl)-α-D-N-acetylneuraminic acid sodium salt hydrate (MU-NANA) substrate. 67 μL of cell supernatant was incubated with 33 µL of MU-NANA (50 µM) in black 96-well black micro-plates. Fluorescence was immediately measured in a kinetic assay over 1 h at Ex = 355 nm and Em = 460 nm.

### Protein-array and ELISA

Protein array and DuoSet ELISA (Human IL-6, and mouse MPO and ALT) were performed according to the manufacturer’s instructions (R&D Systems or Clinisciences for ALT ELISA).

### siRNA transfection

1.25×10^5^ BEAS-2B cells were seeded in a 12-wells plate the day before transfection with specific siRNA or negative control scramble siRNA. Each siRNA stock was diluted to 50 nM in 100 µL of optiMEM (Gibco) containing RNAiMax reagent (ratio siRNA:RNAiMax of 1:3). After 5 min of incubation at room temperature, 100 µL of each siRNA-mix was added to 900 µL of fresh medium per well. Gene knockdown efficacy was evaluated by RT-qPCR after 48 h (medium replaced after 24 h).

### RNA isolation and RT-qPCR

Cells in 6-well plates were lysed with 350 µL RA1 buffer (Macherey-Nagel) and 1/100 diluted β-mercaptoethanol. Total RNA was extracted using the NucleoSpin® RNA kit (Macherey-Nagel), including DNase digestion. RNA concentration was measured with a Nanodrop 2000. cDNA synthesis was performed from 500 ng RNA using the High-Capacity cDNA Reverse Transcription Kit, with IAV M1-specific sense primer or random primers. mRNA levels were quantified by RT-qPCR on a LightCycler 480 (Roche) using 10 ng cDNA, 10 µM primers (M1: sense 5′-AAG ACC AAT CCT GTC ACC TCT GA-3′, antisense 5′-CAA AGC GTC TAC GCT GCA GTC C-3′; CAD: sense 5′-CGT GTT ATT CAG AGG AGC AAG AG-3′, antisense 5′-AGC ATA TGT GGG CGG GAG-3′), and 10 µL SYBR® Premix Ex Taq in a 20 µL reaction volume. Reactions were performed in duplicate, and the thermal protocol included initial denaturation at 95°C for 30 s, followed by 40 cycles of denaturation (95°C for 5 s) and annealing/extension (60°C for 20 s). Melting curves were generated to verify reaction specificity.

### IAV titration by plaque-forming units assay

Titrations in culture media and mouse lungs were performed as previously described [74].

### Transmission electron microscopy

Cells were washed with PBS, detached using trypsin, and centrifuged. They were fixed for 24 h in 4% paraformaldehyde and 1% glutaraldehyde in 0.1 M phosphate buffer (pH 7.2). After washing in PBS, cells were post-fixed with 2% osmium tetroxide for 1 h. Samples were dehydrated in graded ethanol and propylene oxide solutions, then impregnated with a 1:1 mixture of propylene oxide/Epon resin and left overnight in pure resin. The samples were embedded in Epon resin and polymerized at 60°C for 48 h. Ultra-thin sections (90 nm) were cut using a Leica EM UC7 ultramicrotome, stained with 2% uranyl acetate and 5% lead citrate, and analyzed with a JEOL 1011 transmission electron microscope using Digital Micrograph software.

### Scanning electron microscopy

Cells were washed with PBS, detached using trypsin, and centrifuged. They were fixed for 24 hours in 4% paraformaldehyde and 1% glutaraldehyde in 0.1 M phosphate buffer (pH 7.2). After washing in PBS, samples were post-fixed with 2% osmium tetroxide for 1 ho. Samples were dehydrated in a graded ethanol series, then dried in hexamethyldisilazane. The dry samples were placed onto carbon disks and coated with 40 Å of platinum using a GATAN PECS 682 apparatus. Observations were made with a Zeiss Ultra Plus FEG-SEM scanning electron microscope.

### Confocal fluorescence microscopy

BEAS-2B cells were cultured in 12-well plates on cover slides. After treatments, cells were fixed with 4% formaldehyde for 30 min at room temperature and permeabilized with 0.1% Triton X-100 in PBS for 30 min. After blocking with PBS containing 1% BSA and 0.1% Tween 20 for 1 h, cells were stained for 2 h at room temperature with anti-NP-FITC (1/30), anti-NS1 (1/200), and anti-PA (1/50) antibodies. Anti-rabbit-AF488 (2 h at room temperature) served as the secondary antibody for NS1, and anti-mouse-AF488 was used for PA. PBECs were fixed with 4% formaldehyde for 10 min at 4°C, then permeabilized with cold methanol for 10 min at 4°C. After blocking with PBS containing 1% bovine serum albumin and 0.3% Triton X-100 for 10 min, cells were stained for 2 h at room temperature with anti-p63 (1/100), anti-Tubulin (1/100), and anti-Mucin 5AC (1/1000) antibodies. Secondary antibodies used were anti-rabbit-AF546 (for p63), anti-mouse-AF488 (for Mucin 5AC), and anti-mouse-AF647 (for Tubulin). Nuclei were stained with NucBlue reagent for 5 min.

OLC were fixed overnight at 4°C in 4% formaldehyde. Aldehyde groups were quenched with two 10-min incubations in PBS with 0.1% glycine, followed by permeabilization with 0.5% Triton X-100 in PBS for 15 min at room temperature. After 2 h of saturation in PBS with 1% BSA, 0.5% Triton X-100, cells were stained overnight at 4°C with anti-Tubulin (1/500) and anti-NP-FITC (1/50) antibodies. The secondary antibody anti-mouse-AF647 was applied for 2 h at room temperature for Tubulin staining. Nuclei were stained with Hoechst reagent (1/2000) for 10 min. Samples were analyzed and 3D reconstructions were generated using a Leica SP8 confocal microscope with Leica LasX Life Sciences Software.

### Western-blotting

Cells in 6-well plates were lysed with 150 µL of RIPA buffer (with protease inhibitors or PhosphoSafe Extraction Reagent). After centrifugation at 12,000 g for 10 min, protein concentration was measured using the Pierce™ BCA Kit. Ten μg of protein were mixed with Laemmli buffer, heated at 100°C for 5 min, and separated on 12% SDS-PAGE gels. Proteins were transferred to nitrocellulose membranes and probed with primary antibodies: anti-NP (1/500), anti-NS1 (1/1000), anti-PA (1/1000), anti-(P)ERK1/2 (1/2000), anti-(P)AKT (1/1000), anti-(P)p65 (1/3000), or anti-β-actin (1/5000). HRP-conjugated secondary antibodies were used for detection, followed by ECL. Protein bands were visualized using an automated imaging system and analyzed with FUJI FILM MultiGauge software.

### Minigenome assay

The minigenome studies were performed in 24-well plates. Briefly, HEK-293T cells were transfected with together with 50 ng of pRF483-PA-RT, 50 ng of pRF483-PB2-RT, 100 ng of pRF483-NP-RT, 50 ng of pRF483-PB1-RT and 150ng of reporter plasmid pPolI-WSN-NA-firefly luciferase which contains a firefly luciferase ORF flanked by the noncoding regions of the NA segment under the control of human polymerase I promoter. As a negative control, HEK-293T cells were transfected with the same plasmids, with the exception of the PB1 plasmid. The procedure used the Fugene HD transfection reagent according to the manufacturer’s instructions. 20h post – transfection cells were treated with different concentrations of *cis-*aco. 48 h post-transfection, cells were washed twice with PBS and lysed in 100 μl of lysis buffer provided with the Firefly Luciferase Assay System. Firefly luciferase activities were measured on 20 μl of cell extracts, using the Firefly luciferase substrate provided with the above-mentioned kit and a Centro luminometer (Berthold).

### Incucyte® and cell death assays

PBEC were seeded in 96-well plates in the presence of 2 µM of the fluorescent dye SYTOX™, which binds to DNA and rapidly penetrates dying cells upon membrane permeabilization, and then infected with IAV. Real-time cell death assays were performed using an IncuCyte® two-color incubator imaging system. The images obtained were analyzed using the software supplied with the IncuCyte imager, which enables precise analysis of the number of SYTOX™-positive cells present in each image. Experiments were carried out using a minimum of two separate wells for each experimental condition and a minimum of four image fields *per* well.

### Animal infection and fluid collection

7-week-old C57Bl/6 mice (female or male) were intranasally challenged with 10 µg LPS or 200 pfu A/Scotland/20/74 (H3N2) IAV, and treated with 0.6 mg *cis*-aco (30 mg/kg) at various time points. Blood was collected on the sacrifice day, centrifuged at 10,000*g* for 10 min for serum analysis or heparinized for analysis on a ProCyte Dx hematocytometer. Airways were lavaged with 4×0.5 ml PBS for BAL collection, and lungs were perfused with 10 ml PBS injected into the heart. The left lung was fixed in 4% paraformaldehyde for histology. Right lungs were digested enzymatically using the gentleMACS dissociator, and after centrifugation, lung suspensions and BAL fluids were stored at –80°C for subsequent inflammatory mediator analysis. Leukocytes were isolated, red blood cells lysed, and leukocytes counted *via* flow cytometry.

BALB/c NF-κB transgenic mice were used in Ronan Le Goffic’s lab [75]. Mice were challenged with 10 µg LPS or infected with 300 PFU A/Scotland/20/74 (H3N2) IAV. At 1 or 8 days p.i., luciferin (0.75 mg/kg) was administered intranasally, and luciferase activity was measured using the IVIS system.

CAD-deficient C57Bl/6 mice were used in Priscille Brodin and François Trottein’s labs. CAD-deficient and wild-type mice (20 animals *per* group) were infected with 100 PFU A/Scotland/20/74 (H3N2) IAV and treated with 30 mg/kg *cis*-aco 20 min p.i. Body weight loss and survival were monitored daily.

### Genomic DNA Extraction and 16S rRNA Sequencing Analysis

Genomic DNA was extracted from mouse fecal pellets and 16S rRNA sequencing was performed as previously described [76]. The taxonomy of each amplicon sequence variant (ASV) was assigned based on the SILVA database v1.3.8 [77]. ASVs unclassified at the kingdom or phylum level or ASVs classified as Eukaryota or Mitochondria were excluded. Aitchison distances were measured using the microbiome (http://microbiome.github.io) and phyloseq packages [78] in RStudio v4.1.2.

### Flow cytometry analysis

BAL, lungs, or human bronchial epithelial cells were dispensed into round bottomed 96-well plates and centrifuged at 300 *g* at 4°C for 5 min. Samples were further stained using specific antibodies and appropriate isotype controls (listed in Table 3). For each antibody, one well was seeded for the Fluorescence Minus One Control. Dead cells were excluded using the LIVE/DEAD cell staining kit (Invitrogen). Flow cytometry data were acquired on a MACSQuant® Analyzer (Miltenyi Biotec) and analyses were performed using the VenturiOne software (Applied Cytometry).

### Histopathology

Lungs were collected after BAL and airways were washed and placed in 4% paraformaldehyde in PBS. Lung sections of approximately 4 µm thickness were cut and stained with hematoxylin-eosin at the LAPV (Amboise, France). A study pathologist examined the tissue sections using light microscopy on a Leica Diaplan microscope in a blinded experimental protocol. All histopathological findings were graded in a semi-quantitative fashion on a scale of 0 to 4 (0: absent, 1: mild, 2: moderate, 3: severe, 4: extremely severe).

### Cell proliferation and cytotoxicity assays

Cells in 96-well plates were washed twice with PBS and incubated for 1 h at 37°C with 100 µL of MTS reagent diluted 1/5 for the cell proliferation test. Optical density was measured at 490 nm. Cells were stained for 15 min at 4°C with Live/Dead (1/1000^e^), anti-Ki67(1/100^e^) or MitoTracker(1/5000^e^), or 5 min at 37°C with Dihydrorhodamine(DHR)-123(1/100^e^) before flow cytometry analysis.

### Statistical analyses

Statistical analyses were conducted using GraphPad Prism on raw data. Data are presented as mean ± SEM. The statistical values, including the number of replicates (n) and the statistical test used, are detailed in the figure legends. *p < 0.05, **p < 0.005, ***p < 0.0005, ****p < 0.0001. For *in vitro* experiments, “n” refers to the number of separate experiments, while for *in vivo* studies, “n” refers to the number of individual animals.

## DATA AVAILABILITY STATEMENT

The datasets generated during and/or analysed during the current study are available from the corresponding author on reasonable request.

## CONTRIBUTIONS DATA

**AC** Data curation, Investigation, Conceptualization, Formal analysis, Supervision, Validation, Visualization, Methodology, Writing – original draft; Writing – review & editing. **VV, AW, DD** Data curation, Investigation, Formal analysis, Validation, review & editing, Methodology. **LG, SH** Investigation, Methodology. **LC, FJ, CM, LPC** Investigation, Methodology, Resources, Writing – review & editing. **RLG** Resources, Investigation, Validation, Writing – review & editing. **BDC** Resources, Investigation, Validation. **AM, EH, FT** Resources, Validation, Methodology. **JBG, DT, MGS** Resources, Investigation, Formal analysis, Validation. **TB** Investigation, Methodology, Validation, review & editing. **DF** Investigation. **CP** Formal analysis, Resources, Validation; review & editing, review & editing. **AG** Supervision, Validation, Writing – review & editing. **MS-T** Conceptualization, Formal analysis, Resources, Validation, Supervision, Funding acquisition, Supervision, Project administration, Writing – review & editing.

## ACKNOWLEDGMENTS

The authors are grateful to Pr. Pieter Hiemstra (Laboratory for Respiratory Cell Biology and Immunology of the Department of Pulmonology of the Leiden University Medical Center (LUMC)) for his collaboration in setting up the *ex vivo* culture model of primary human lung epithelial cells. Also, we thank Benoit Briard and Sandra Khau (CEPR, Inserm, Tours) for their advice with the Incucyte® and cell death assays.

## EXPANDED VIEW CONTENT

**Figure EV1.** Tolerance of bronchial epithelial cells to metabolite exposure. **(a)** BEAS-2B cells were treated with medium alone or with 3.4 mM *cis*-aco (CA), itaconate (Ita), oxaloacetate (Oxa), isocitrate (IsoC), citrate (Cit), fumarate (Fum), pyruvate (Pyr) or glucose (Glc) for 16h. Cytotoxicity was assessed using the MTS assay. **(b-e)** BEAS-2B cells were treated (blue) or not (white) with 1.2 or 2.3 or 3.4 mM *cis*-aco for 20 h **(b)** or with 3.4 mM *cis*-aco for 6 or 24h **(c-e)**. Effects on cell viability **(b)**, proliferation **(c)**, mitochondrial labeling **(d)**, and ROS production **(d)** were analyzed using Live Dead, Ki67, mitotracker and DHR123 staining respectively. **(f)** PBEC were treated with 2.3, 3.4 or 5.7 mM *cis*-aco for 16h, and cytotoxicity was assessed using the MTS assay. Data are presented as the mean ± SEM from 4 **(b-f**) or 5 **(a)** independent experiments. Statistical analyses were performed using the Kruskal-Wallis test with Dunn’s multiple comparison test **(a-c)** or the Friedman test with Dunn’s multiple comparison test **(d-f)**.

**Figure EV2.** Comparison of anti-influenza effects of *cis*-aco and *trans*-aco. BEAS-2B cells were incubated with medium alone or with increasing concentrations of *cis*-aconitic (CA) or *trans-*aconitic acid (TA) for 20 h. The chemical structures of these metabolites are shown in **(a)**. **(b)** Cell viability was assessed using the MTS assay. A viability threshold of 80% was set (red dashed line). **(c, d)** BEAS-2B cells were infected with influenza A/Scotland/20/74 (H3N2) virus (IAV) at an MOI=1 (pink bars) or left uninfected (NI) for 4 h, followed by treatment or not with increasing concentrations of CA (blue) or TA (green) for 16h. **(c)** Viral particle production was assessed using a neuraminidase activity assay. **(d)** IL-6 levels in cell supernatants were measured by ELISA. Data are presented as the mean ± SEM and represent cumulative results from 4 or 5 independent experiments. Statistical analysis was performed using the Kruskal-Wallis test with Dunn’s multiple comparison test.

**Figure EV3.** *Cis*-aco protects against IAV-induced cell death in human primary bronchial epithelial cells (PBEC). (**a-b**) PBEC were infected with virus A/Scotland/20/74 (H3N2) at an MOI=1, and treated or not with 3.4 mM *cis*-aco (CA) 10 min p.i.. SYTOX^TM^ labelling was monitored over 18h. Representative images (**(a)**; scale bar: 200 µm) and quantification of labeling **(b)** at 18 h p.i. were used to assess cell death. Data are presented as the mean ± SEM from duplicate PBEC derived from 4 independent patients analyzed. Statistical analysis was performed using the Friedman test with Dunn’s multiple comparison test.

**Figure EV4.** Tolerance and safety of *cis*-aco *in vivo*. Seven-week-old female and male C57Bl/6 mice were intranasally instilled with PBS or 30 mg/kg *cis*-aco (CA) for 15 days, according to the schedule outlined. (**a)** Body weight was monitored throughout the study. **(b)** Mice were euthanized on day 15 and serum was collected to determine ALAT activity levels. **(c)** Microbiota alterations were assessed in mouse fecal pellets, after genomic DNA extraction and 16S rRNA sequencing. Microbial community diversity was quantified using Aitchison distance values, a measure of beta diversity. Each point represents the Aitchison distance between a CA-treated mouse and a PBS-treated mouse at day 0, 7, or 14. **(d)** The number of immune cells in BAL fluids was assessed by flow cytometry. **(e)** Blood cell analysis was also conducted. Data from **(e)** represent pooled results from 3 mice. All other data are presented as the mean ± SEM and are cumulative **(a-c)** or representative (**d)** of 3 independent experiments. Statistical analysis was performed using the Kruskal-Wallis test with Dunn’s multiple comparison test.

**Figure EV5.** Confirmation of the protective effects of *cis*-aco in IAV-infected Balb/c mice. To validate the findings observed in C57Bl/6 mice (see Figures 3 a-d) in a different mouse strain and sex, we conducted an experiment using male Balb/c mice. These (NF-κB transgenic) Balb/c animals were infected with 300 PFU of A/Scotland/20/74 (H3N2) IAV and treated intranasally with 30 mg/kg *cis*-aco (CA) 2 days p.i.. Survival **(a)** and body weight **(b)** were monitored daily. All data are presented as the mean ± SEM. Statistical analysis was performed using the Log-rank (Mantel-Cox) test.

