## Supplementary material for "*Cis*-aconitate therapy protects against influenza mortality by dual targeting of viral polymerase and ERK/AKT/NF-κB signaling": Tables 1-3

**TABLE 1 : Patients’ characteristics for OLC preparations**

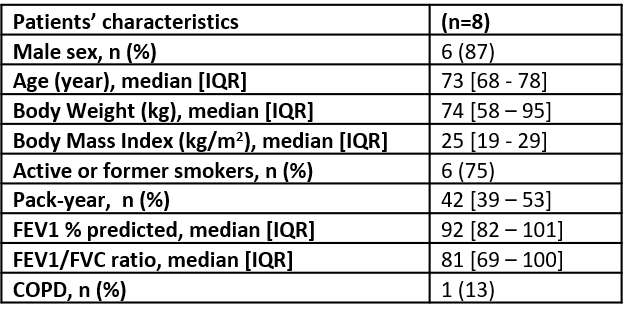

IQR: interquartile range ; FEV1 :forced expiratory volume in 1 second ; FVC : forced vital capacity; COPD : chronic obstructive pulmonary disease.

**TABLE 2. Characteristics of patients hospitalized for community-acquired pneumonia (CAP).** Quantitative data are reported as the median value and interquartile range [IQR] and qualitative value are reported as n (%).

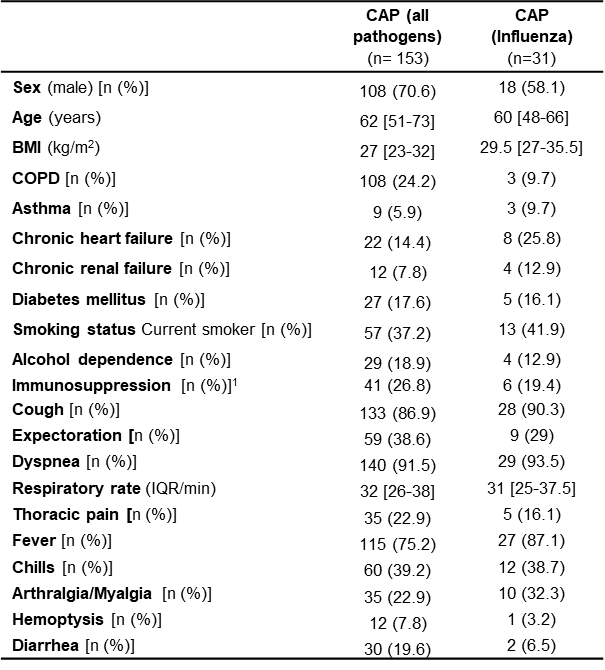

*^1^ defined as solid cancer, hemopathy, organ transplant, bone marrow transplant, HIV infection, and splenectomy*

**TABLE 3: List of reagents and resources used**

| **REAGENT OR RESOURCE** | **SOURCE** | **IDENTIFIER** |
| --- | --- | --- |
| **Antibodies** |  |  |
| Anti-influenza A Virus Nucleoprotein antibody | Abcam | ab128193 |
| Anti-influenza A Virus Nucleoprotein antibody FITC | Abcam | ab20921 |
| Anti-Influenza A PA antibody | Invitrogen | PA532223 |
| Anti-Influenza NS1 antibody | Gift from Dr Marc, INRAE, Nouzilly, France | N/A |
| beta Actin Monoclonal Antibody | ThermoFisher Scientific | MA5-15739 |
| Anti-Phospho-Akt (Ser473) | Cell Signaling Technology | 4060 |
| Anti-Phospho-ERK1/2 | Cell Signaling Technology | 4377 |
| Phospho-NFκB p65 (Ser536) Monoclonal Antibody (T.849.2) | ThermoFisher Scientific | MA5-15160 |
| Anti-Mouse IgG (whole molecule)–Peroxidase antibody | Sigma-Aldrich | A9044 |
| Anti-Rabbit IgG (whole molecule)–Peroxidase antibody | Sigma-Aldrich | A9169 |
| APC-eFluor780-conjugated anti-CD45 (30-F11) | ThermoFisher Scientific | 47-0451-82 |
| CD86 (B7-2) Monoclonal Antibody (GL1), FITC | eBiosciences | 11-0862-82 |
| MHC Class II (I-A) Monoclonal Antibody (NIMR-4), PE | eBiosciences | 12-5322-81 |
| CD11b Monoclonal Antibody (M1/70), PerCP-Blueine5.5 | eBiosciences | 45-0112-82 |
| CD335 (NKp46) Monoclonal Antibody (29A1.4), eFluor 450 | eBiosciences | 48-3351-82 |
| APC Rat Anti-Mouse Ly-6G antibody | BD Biosciences | 560599 |
| PE/Blueine7 anti-mouse CD11c Antibody | BioLegend | 117318 |
| CD8a Monoclonal Antibody (53-6.7), eFluor 450 | eBiosciences | 48-0081-82 |
| PE/Blueine7 anti-mouse/human CD11b Antibody | BioLegend | 101216 |
| FITC anti-mouse CD4 Antibody | BioLegend | 130308 |
| BB700 Mouse Anti-Mouse NK-1.1 Antibody | eBiosciences | 566502 |
| CD69-APC, mouse Antibody | Miltenyi | 130-115-576 |
| PE/Blueine7 anti-mouse CD3ε Antibody | BioLegend | 100320 |
| CD3e Monoclonal Antibody (145-2C11), PE | eBiosciences | 12-0031-83 |
| F4/80 Monoclonal Antibody (BM8), eFluor 660 | eBiosciences | 50-4801-82 |
| Dihydrorhodamine 123 | Sigma-Aldrich | 109244-58-8 |
| MitoTracker™ Red CM-H2Xros | ThermoFisher Scientific | M7513 |
| V450 Mouse anti-Ki-67 | BD Biosciences | 561281 |
| Anti-p63 Antibody | Abcam | 124762 |
| Anti-tubulin Antibody | Sigma | T6793 |
| Anti-Mucin5Ac | ThermoFisher Scientific | MA5-12178 |
| Goat anti-rabbit antibody AF 546 | ThermoFisher Scientific | A11035 |
| Goat anti-mouse antibody AF 488 | ThermoFisher Scientific | A21121 |
| Goat anti-mouse antibody AF 647 | ThermoFisher Scientific | A21242 |
| **Chemicals, peptides** |  |  |
| Sodium Pyruvate (100 mM) (Gibco™) | ThermoFisher Scientific | 11360070 |
| Lipopolysaccharide from Escherichia coli 0111:B4 | Invivogen | tlrl-eblps |
| DL-Isocitric acid trisodium salt hydrate | Sigma-Aldrich | I1252 |
| Triethyl citrate | Sigma-Aldrich | 14849 |
| Sodium fumarate dibasic | Sigma-Aldrich | F1506 |
| D-(+)-Glucose | Sigma-Aldrich | G7021 |
| Itaconic acid, +99 %, ACROS Organics™ | Fisher Scientific | 10457700 |
| Oxaloacetic acid | Sigma-Aldrich | O4126 |
| cis-aconitic acid | Sigma-Aldrich | A3412 |
| trans-aconitic acid | Sigma-Aldrich | 122750 |
| Poly(I:C) LMW 25mg | Invivogen | tlrl-picw |
| Phorbol 12-myristate 13-acetate | Sigma-Aldrich | 16561-29-8 |
| Recombinant Human TNF-alpha Protein | R&D systems | 210-TA |
| ActinRed™ 555 ReadyProbes™ Reagent | ThermoFisher Scientific | R37112 |
| NucBlue™ Fixed Cell ReadyProbes™ Reagent | ThermoFisher Scientific | R37606 |
| 2′-(4-Methylumbelliferyl)-α-D-N-acetylneuraminic acid sodium salt hydrate | Chemodex | M0096 |
| Protease Inhibitor Cocktail | Sigma Aldrich | P8340 |
| PhosphoSafe Extraction Reagent | Sigma Aldrich | 71296 |
| Gibco^TM^ Ham's F-12 Nutrient Mix | ThermoFisher Scientific | 31765027 |
| Gibco^TM^ MEM | ThermoFisher Scientific | 31095029 |
| Gibco™ GlutaMAX™ Supplement | ThermoFisher Scientific | 13462629 |
| Gibco™ HEPES (1M) | ThermoFisher Scientific | 11560496 |
| BEGM™ Bronchial Epithelial Cell Growth Medium BulletKit™ | Lonza | CC-3170 |
| Trypsin 0.25 %/EDTA 0.02 % in PBS | PAN BIOTECH | P10-020100 |
| Trypsin, TPCK Treated | ThermoFisher Scientific | 20233 |
| Trypsin / Lys-C Mix, Mass Spec Grade | Promega | V5072 |
| MEM Eagle with Earle's BSS (2X) | Lonza | BE12-668F |
| Crystal Violet Oxalate | RAL Diagnostics | 361490 |
| Formaldehyde, 37 wt % sol. in water, stab. with 5-15% methanol | Acros Organics | 119690010 |
| Avicel® RC 581 Stabilizer | FMC BioPolymer | N/A |
| Annexin V-FITC kit | Miltenyi Biotech | 130-092-052 |
| SYTOX™ Green nucleic acid stain | InVitroGen | S7020 |
| True-Nuclear™ Transcription Factor Buffer Set | Biolegend | 424401 |
| BD Cytofix/Cytoperm™ Fixation/Permeabilization Solution Kit | Fisherscientific | BDB554714 |
| Red Blood Cell Lysing Buffer Hybri-Max™ | Sigma Aldrich | R7757 |
| TB Green® Premix Ex Taq™ | Takara | RR420L |
| 50% EM Glutaraldehyde | TAAB Laboratory Equipment | G045 |
| Uranyl acetate | Merck | 8473 |
| Osmium tetroxide 4% solution | Electron Microscopy Science | 19150 |
| Oseltamivir phosphate | Sigma Aldrich | 204255-11-8 |
| Gibco™ optiMEM | Fischer Scientific | 31985070 |
| Invitrogen™ Lipofectamine™ RNAiMAX Transfection Reagent | Fischer scientific | 13-778-150 |
| ON-TARGETPlus human smartpool IRG1 | Dharmacon | L-180668-01-0005 |
| MISSION® negative control scramble | Sigma Aldrich | SIC001 |
| Gibco™ Milieu Hibernate™-A | Fischer Scientific | 12087586 |
| Primocin | Invivogen | ant-pm-05 |
| Bovine Albumin Fraction V | Fischer Scientific | 15260037 |
| PureCol | Advanced BioMatrix | 5005-B |
| Human Fibronectin Stabilized Solution | PromoCell | C-43060 |
| Heparin | StemCell | 07980 |
| Hydrocortisone | StemCell | 07925 |
| Soybean Trypsin Inhibitor | Sigma Aldrich | T-9128 |
| Isoproterenol | Sigma Aldrich | I-6504 |
| Bovine pituitary extract | Fischer Scientific | 11568866 |
| Epidermal Growth Factor (EGF) | Fischer Scientific | 10134762 |
| Serum-free keratinocyte medium (Gibco) | Fischer Scientific | 11590526 |
| Ca^2+/^Mg^2+^-free Hank’s Balanced Salt Solution | Gibco | 88284 |
| Proteinase type XIV | Sigma Aldrich | P5147-100MG |
| Hexamethyldisilazane | Sigma Aldrich | 440191-100ML |
| GATAN PECS 682 apparatus | Pleasanton, CA | https://www.gatan.com |
| Fugene HD transfection reagent | Promega | E2311 |
| Firefly Luciferase Assay System | Promega | E1500 |
| fluorescent dye SYTOX™ | Fischer scientific | 10768273 |
| PneumaCult™­Ex Kit | StemCell | 05001 |
| PneumaCult™­Ali Kit | StemCell | 05008 |
| **Commercial Assays** |  |  |
| LIVE/DEAD™ Fixable Aqua Dead Cell Stain Kit | ThermoFisher Scientific | L34966 |

| Phusion™ High-fidelity DNA polymerase | ThermoFisher Scientific | 16237911 |
| --- | --- | --- |

| Pierce™ BCA Protein Assay Kit | ThermoFisher Scientific | 23225 |
| --- | --- | --- |
| CellTiter 96® AQueous One Solution Cell Proliferation Assay | Promega | G3582 |
| Nucleospin® RNA | Macherey-Nagel | 740955 |
| High Capacity cDNA reverse transcription kit | Applied Biosystems | 4368813 |
| Human IL6 ELISA DuoSet | R&D Systems | DY206 |
| Mouse MPO ELISA DuoSet | R&D Systems | DY3667 |
| Human Cytokine Array Kit | R&D Systems | ARY005B |
| Mouse XL Cytokine Array | R&D Systems | ARY028 |
| SequalPrep normalization kit | Thermofisher | A1051001 |
| MagMAX™ DNA Multi-Sample Kit | Thermofisher | 4413020 |
| **Experimental Models** |  |  |
| C57Bl/6 mice | Janvier | C57BL/6JRjFEMELLESPF4 |
| CAD-deficient C57Bl/6 mice | Dr. Priscille BRODIN (University of Lille, France) | N/A |
| BALB/c NF-kB transgenic mice | Dr. Ronan LE GOFFIC  (INRAE, VIM, Jouy-en-Josas, France) | N/A |
| BEAS-2B | ATCC® | CRL-9609 |
| A549 | ATCC® | CCL-185 |
| MDCK.2 | ATCC® | CRL-2935 |
| HEK293T | ATCC® | CRL-11268 |
| B/Paris/234/2013 (Yamagata) lineage | European Virus Archive Global (EVAg). | 014V-01887 |
| Influenza A/Scotland/20/74 | Pasteur Institute, France | N/A |
| Influenza A/PR/8/34 (PR8) | kindly provided by Dr. Georg Kochs (Freiburg University, Germany) | N/A |
| Influenza H1N1 pdm09 | generously given by Dr. François Trottein (Center for Infection and Immunity of Lille) | N/A |
| **Softwares** |  |  |
| GraphPad Prism | GraphPad Software | https://www.graphpad.com/scientific-software/prism/ |
| VenturiOne | Applied Cytometry | https://www.appliedcytometry.com/venturi/ |
| FUJI FILM Multigauge | Bioz | https://www.bioz.com/ |
| LightCycler 480 SW V.1.5 | Roche | https://lifescience.roche.com/ |
| ImageJ | Imagej | https://imagej.net/Welcome |
| BioStation IM software (v2.12) | Nikon | https://www.nikon.com/products/microscope-solutions/ |
| Leica LasX Life Sciences | Leica Microsystems | https://www.leica-microsystems.com |
| Digital Micrograph V.3 | Gatan | https://www.gatan.com/products/tem-analysis |
| MagMAX Express 96-Deep Well Magnetic Particle Processor | Applied Biosystems | 4472991 |
| gentleMACS dissociator | Miltenyi Biotec | 130-093-235 |
| ProCyte Dx hematocytometer | Idexx | https://www.idexx.fr |
| IncuCyte® two-color incubator imaging system | Essen Biosciences, Sartorius | https://www.essenbioscience.com |
| MF ChemiBis 3.2, | DNR BioImaging Systems | https://hvdlifesciences.at/dnr-bio-imaging-systems.html |
| Zeiss Ultra plus FEG-SEM scanning electron microscope | Zeiss | https://www.zeiss.com |
