## Supplementary figures 1 – 5 for "*Cis*-aconitate therapy protects against influenza mortality by dual targeting of viral polymerase and ERK/AKT/NF-κB signaling"

### Slide 1
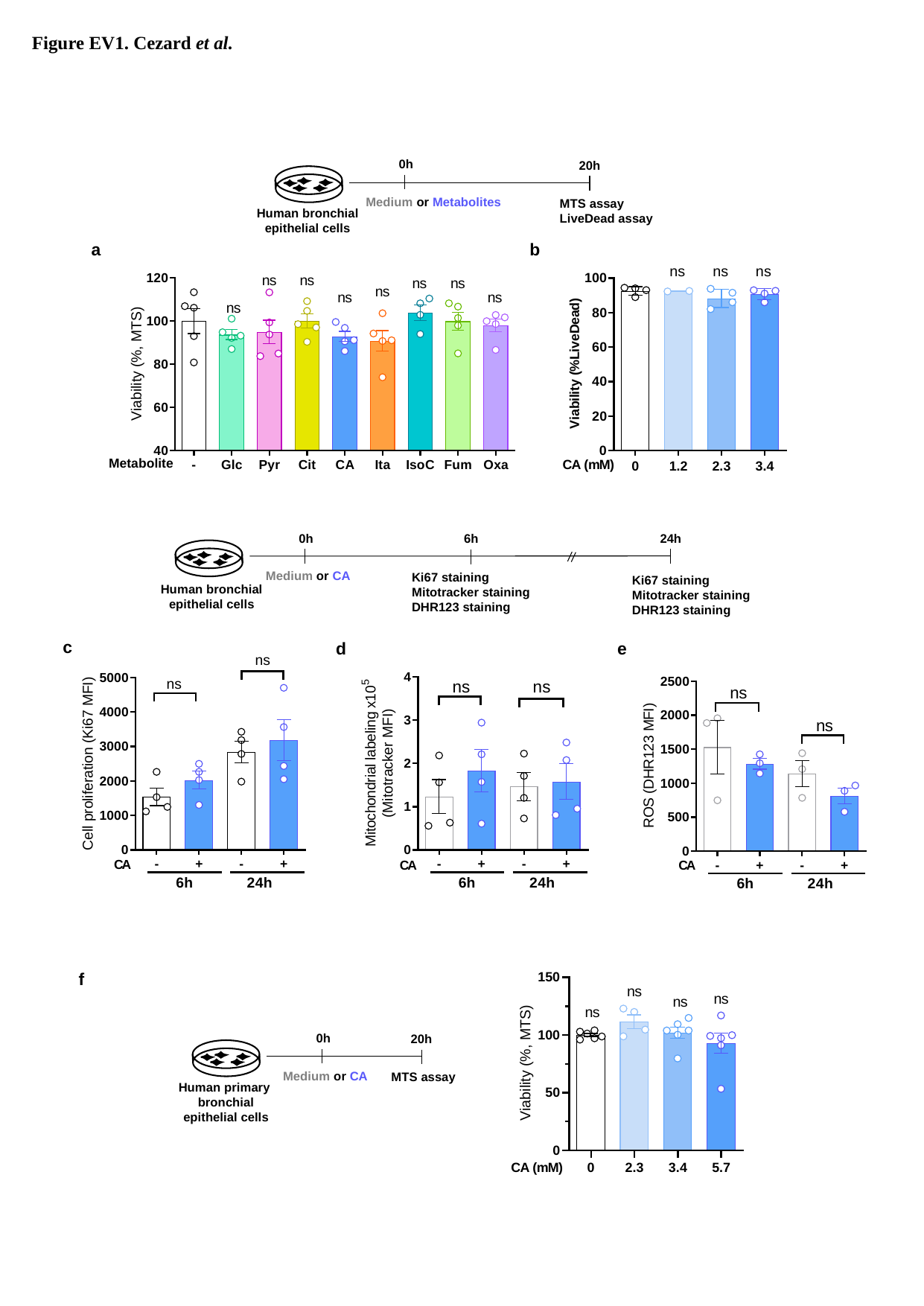

Figure EV1. Cezard et al.
0h
20h
Medium or Metabolites
MTS assay
LiveDead assay
Human bronchial epithelial cells
a
b
0h
24h
6h
Medium or CA
Ki67 staining
Mitotracker staining
DHR123 staining
Ki67 staining
Mitotracker staining
DHR123 staining
Human bronchial epithelial cells
c
d
e
f
0h
20h
Medium or CA
MTS assay
Human primary bronchial epithelial cells

### Slide 2
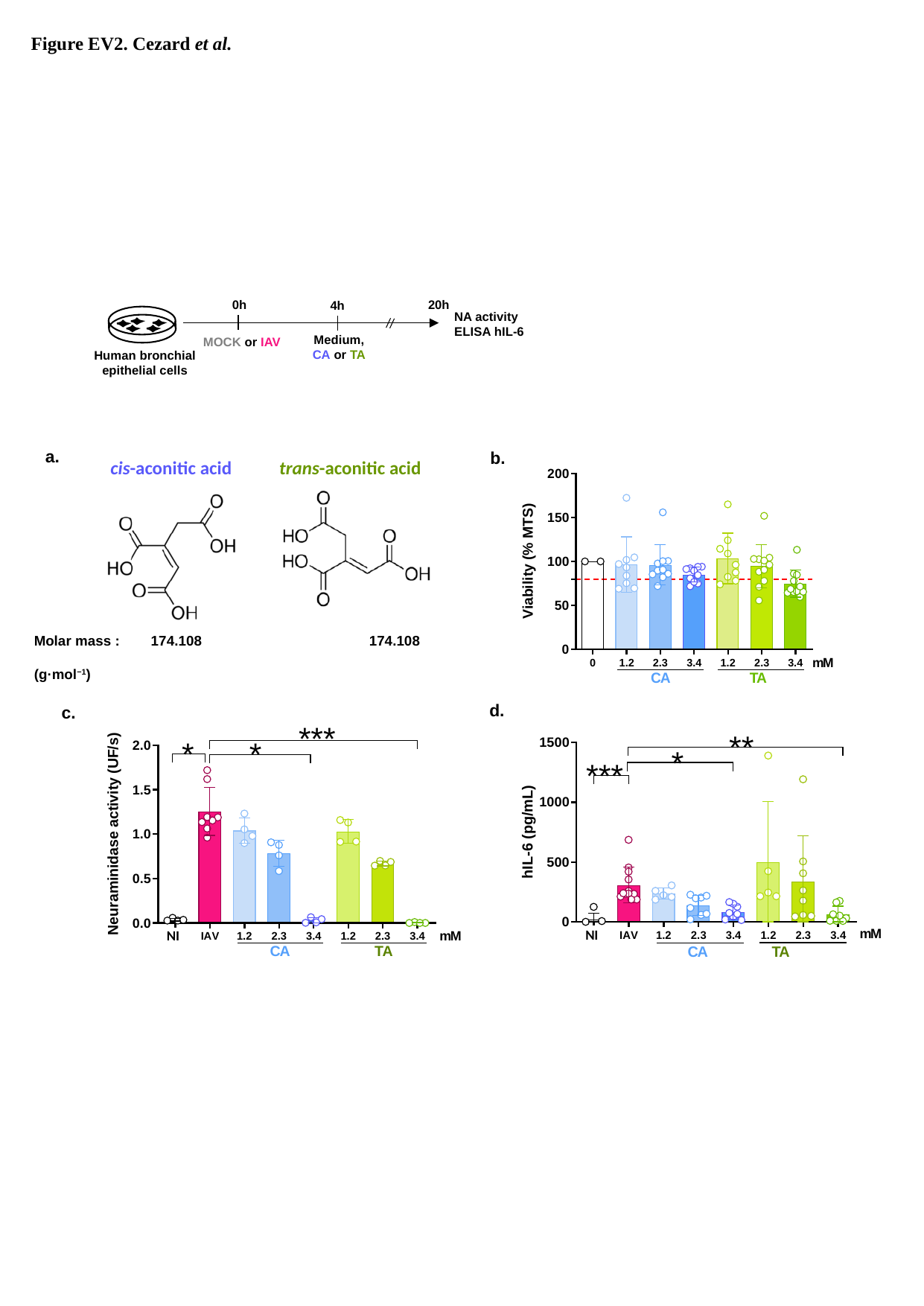

Figure EV2. Cezard et al.
0h
20h
4h
NA activity
ELISA hIL-6
Medium, CA or TA
MOCK or IAV
Human bronchial epithelial cells
a.
b.
cis-aconitic acid
trans-aconitic acid
Molar mass : 174.108 	 	174.108
(g·mol^−1^)
d.
c.
NI
NI

### Slide 3
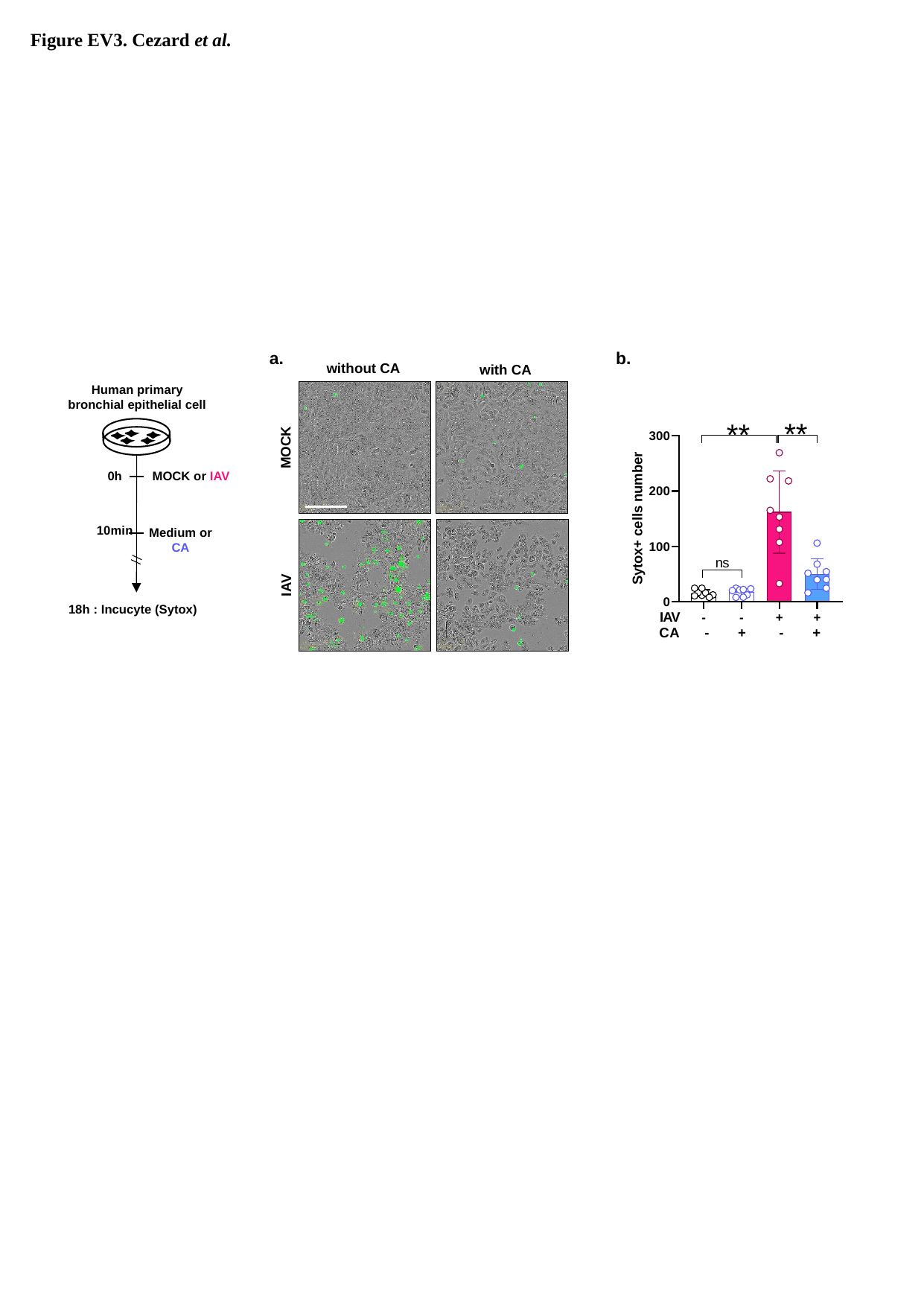

Figure EV3. Cezard et al.
a.
b.
without CA
with CA
MOCK
IAV
Human primary bronchial epithelial cell
0h
MOCK or IAV
10min
Medium or CA
18h : Incucyte (Sytox)

### Slide 4
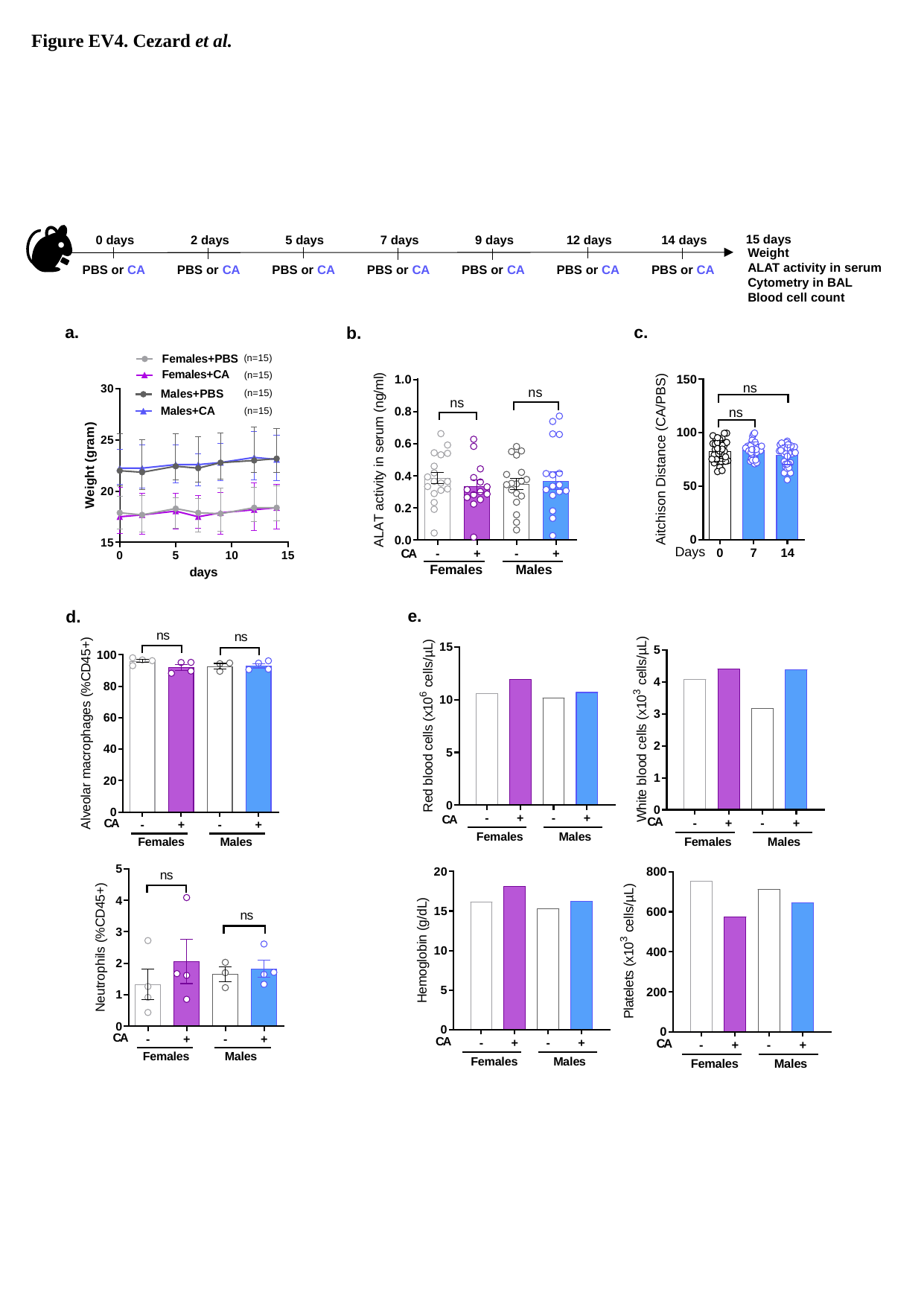

Figure EV4. Cezard et al.
15 days
0 days
2 days
5 days
7 days
9 days
12 days
14 days
Weight
ALAT activity in serum
Cytometry in BAL
Blood cell count
PBS or CA
PBS or CA
PBS or CA
PBS or CA
PBS or CA
PBS or CA
PBS or CA
a.
c.
b.
e.
d.

### Slide 5
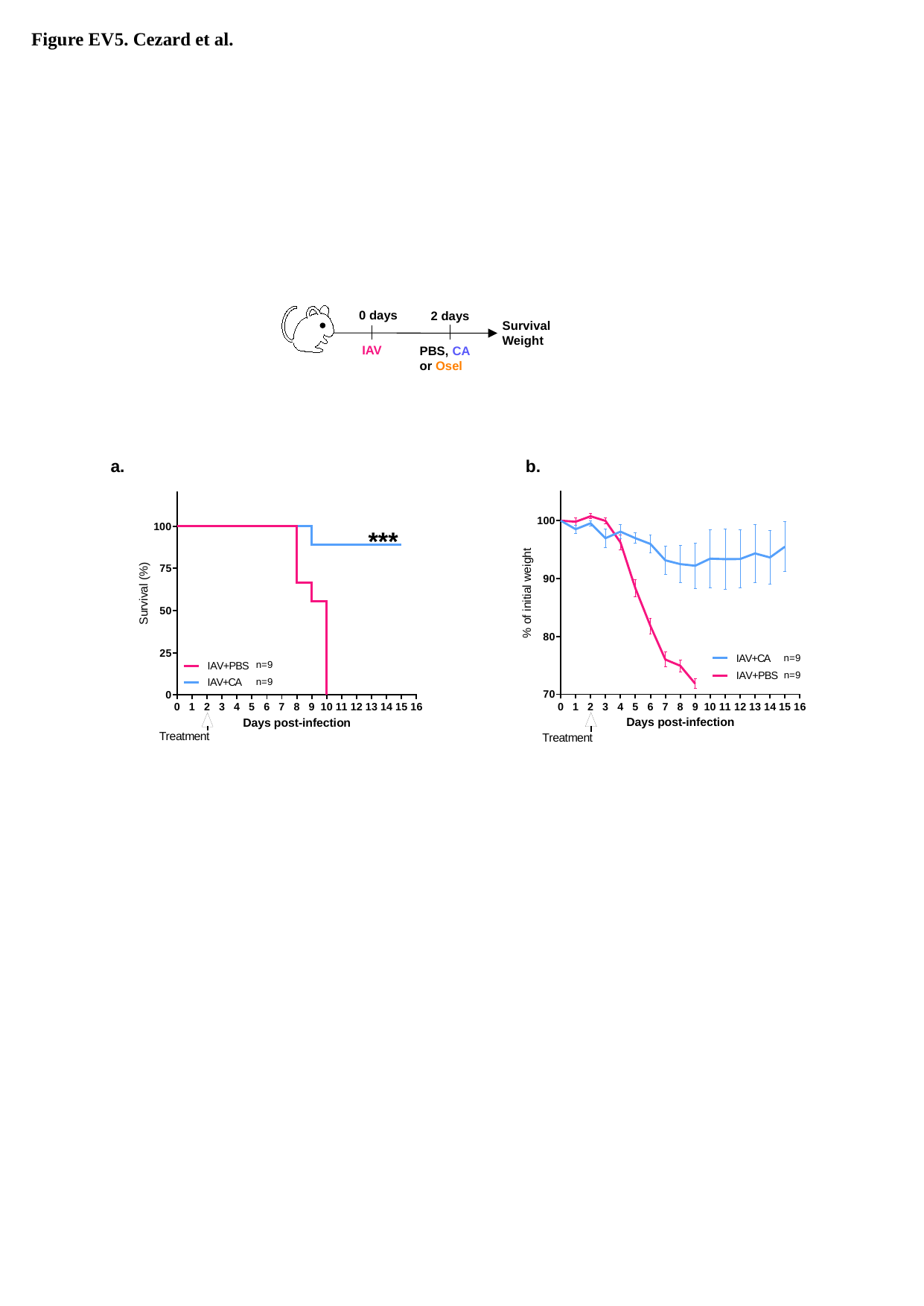

Figure EV5. Cezard et al.
0 days
2 days
Survival
Weight
IAV
PBS, CA or Osel
a.
b.
